# Environment-Aware DNA Language Model for Stress-Responsive Genomic Prioritization in Maize

**DOI:** 10.64898/2026.09.11.749989

**Authors:** Debasmita Pal, Aaron Odell, Anuradha Singh, Addie Thompson, Arun Ross, Anne Thessen

## Abstract

Abiotic stresses such as heat and drought severely reduce the productivity of major crops including maize, with adverse implications for agricultural sustainability. Despite progress in characterizing stress resilience, identifying genomic regions associated with stress response remains challenging due to complex genotype-by-environment interactions. Genomic Foundation Models (GFMs), inspired by advances in Large Language Models (LLMs), leverage large-scale self-supervised pre-training on DNA sequences to capture regulatory patterns; however, their adaptation to plant stress-response analysis remains underexplored. In this work, we propose an environment-aware DNA-LLM framework that incorporates stress-specific prompt tokens into AgroNT, a transformer-based plant GFM, using parameter-efficient fine-tuning (PEFT). The model learns stress-conditioned genomic representations that exhibit distinct structures across environmental contexts. We further formulate a prioritization method that uses stress-induced shifts in these representations relative to the control condition together with stress-token-associated transformer attention patterns to identify putative stress-responsive genomic regions. The framework was evaluated on the Genomes-to-Fields (G2F) dataset, comprising over 4,000 maize hybrids grown across 38 locations over 8 years, using grain yield under heat, drought, and combined heat-drought stress. Flowering traits were used to assess generalizability of the stress-conditioned representations across phenotypes. The prioritized regions were assessed using spatiotemporal differential gene-expression profiles, stress-associated quantitative trait loci, and transcription-factor characterization. Regulatory motif analysis further revealed stress-associated motifs enriched within the prioritized sequences as well as model-emphasized high-attention regions. Functional annotation of genes proximal to prioritized regions highlighted roles in transcriptional regulation, signaling, and metabolic pathways relevant to abiotic stress adaptation. Collectively, these findings demonstrate the potential of stress-conditioned transformer-based sequence modeling for environment-aware genome-to-phenome analysis in crop improvement.

## 1. Introduction

Abiotic stresses such as heat and drought severely impair plant growth and development, leading to substantial reductions in crop productivity [1]. Ongoing climate change is exacerbating these challenges by increasing the duration, frequency and intensity of such stresses, posing serious threats to global food security [2–4]. Maize (Zea mays), one of the most widely cultivated staple crops and a major source of food, animal feed, and biofuel, is particularly vulnerable to adverse environmental conditions [5,6]. Therefore, elucidating the genomic basis of stress adaptation is essential for developing stress-tolerant cultivars and sustaining agricultural productivity [7]. Despite progress in characterizing stress response using Genome-Wide Association Studies (GWAS), identifying stress-responsive genomic regions remains a challenge because stress resilience is a complex, polygenic trait influenced by genotype-by-environment (G×E) interactions [8,9]. Recent advances in Artificial Intelligence (AI) offer new opportunities for extracting intricate patterns from biological data [10].

In particular, Large Language Models (LLMs) have driven a paradigm shift through large-scale self-supervised pre-training on massive text corpora, enabling a range of natural language processing tasks, such as text generation, summarization, and question answering [11]. These models use neural network architectures, specifically transformers [12], which utilize attention mechanisms to capture long-range dependencies across input tokens. Tokens (e.g., words or subwords) are mapped into a high-dimensional embedding space where each token is represented as a numerical vector learned during pre-training. These representations encode contextual relationships among tokens, with semantically related representations often exhibiting closer proximity in the learned embedding space.

Building on the success of LLMs, similar representation-learning strategies have been extended to biological sequences, giving rise to DNA-LLMs, also referred to as Genomic Foundation Models (GFMs) [13,14]. Through *large-scale self-supervised pre-training* on *unlabeled nucleotide sequences,* often derived from reference genomes, pan-genomes, or multi-species genomic datasets, these models learn representation of sequence context and regulatory patterns, providing a promising approach for deciphering DNA sequences, particularly non-coding regions [15]. GFMs can subsequently be fine-tuned for various tasks, including regulatory element classification, DNA methylation prediction, variant effect prediction, gene expression modeling, and *in silico* sequence design [14,16].

In this study, we introduce an *environment-aware* DNA-LLM framework to identify stress-responsive genomic regions associated with maize grain yield under heat, drought, and combined heat-drought conditions. Flowering traits were additionally used to assess cross-phenotype generalizability of the learned representations. We incorporated stress-specific special tokens as prompts into the Agronomic Nucleotide Transformer (AgroNT) [17], a transformer-based GFM pre-trained primarily on crops and other plant genomes. Through parameter-efficient fine-tuning (PEFT) [18], the model learned to encode *stress-conditioned genomic representations.* We further formulated a prioritization method that leverages the stress-induced shifts relative to control conditions in the learned sequence embedding space together with stress-token-associated attention patterns to identify putative regulatory regions relevant to stress adaptation. We used the Genome to Fields (G2F)^1^ dataset curated by Lopez-Cruz et al. [19], comprising phenotypic measurements from over 4,000 maize hybrids evaluated under diverse environmental conditions. To our knowledge, this work represents the first application of a GFM to plant stress response analysis through explicit modeling of stress-conditioned sequence context.

Traditionally, GWAS [20] is widely used to identify genetic variants, typically Single Nucleotide Polymorphisms (SNPs), associated with plant responses across environmental conditions [8,9,21], providing a powerful tool for candidate-locus discovery. However, GWAS results are sensitive to the choice of significance thresholds, which influences the balance between false-positive and false-negative findings [22]. Candidate prioritization therefore often relies on post-GWAS strategies, such as functional annotation, pathway enrichment, meta-analysis, and expression quantitative trait loci (eQTL) integration [23]. Classical machine learning methods, including Random Forests, Support Vector Machines, have also been explored to analyze genetic variation and prioritize QTLs and candidate genes. Nevertheless, these methods typically depend on engineered features or summary statistics and do not explicitly model the underlying sequence context [24–26]. Deep learning models such as DeepSEA [27], PlantDeepSEA [28], Enformer [29], have demonstrated the ability to learn informative representations directly from biological sequences for predicting regulatory activity and variant effects. However, these supervised approaches remain constrained by the availability and quality of annotated training data [25].

In contrast, self-supervised pre-training adopted by GFM allows models to learn latent sequence patterns from large-scale unlabeled genomic datasets and generate representations that can subsequently be adapted to diverse downstream tasks [14]. In this work, we leveraged this capability by integrating transformer-based sequence modeling with stress-conditioned representations to encode the genomic context surrounding candidate SNPs identified by GWAS at a permissive significance threshold. Because the sequence regions flanking these loci may contain regulatory signals associated with gene function and stress responses, the learned representations provide an environment-aware basis for prioritizing putative stress-responsive genomic regions. The prioritized loci were compared with GWAS-based rankings to assess whether the model-based prioritization captured information beyond conventional statistical association. Biological relevance was further assessed through complementary evidence from transcriptomic overlap, stress-associated meta-QTL (MQTL) analysis, transcription-factor (TF) characterization, regulatory motif analysis, and functional annotations of proximal genes. Source code is available on GitHub^2^.

## 2. Materials and Methods

### 2.1. Data Preparation and GWAS

We utilized the curated G2F genotypic and phenotypic datasets from Lopez-Cruz et al. [19], generated through an automated pipeline that integrates publicly available weather and soil data and derives environmental covariates (ECs) using APSIM crop model [30]. Heat-and drought-stress indices were also computed from these ECs over the reproductive period from flag-leaf emergence to grain filling. These included the water-supply demand ratio (SDR) and HI30, the number of days with a maximum temperature exceeding 30°C. The dataset comprises 78,686 phenotypic observations for grain yield (ton/ha) and flowering traits, including days to anthesis, days to silking, anthesis-silking interval (ASI), from 4,372 hybrids evaluated over 8 years (2014-2021) across 38 locations, representing 136 year-location combinations. Based on the stress indices, these 136 combinations were categorized into four environmental conditions: *no stress, heat stress, drought stress and combined heat-drought stress*, summarized in Table 1. After filtering for minor allele frequency (MAF) and missingness, followed by linkage disequilibrium (LD) pruning, the genotype panel contained 98,026 SNPs mapped to the B73 reference genome (Zm-B73-REFERENCE-NAM-5.0).

**Table 1.** Summary of G2F datasets used in GWAS, curated by Lopez-Cruz et al. [. **19]**

| **Stress Condition** | **Stress Indices** | **No. of Hybrids Tested** | **No. of Year-Locations** |
|---|---|---|---|
| No Stress (Control) | HI30 <= 36 and SDR > 0.54 | 4,298 | 80 |
| Heat | HI30 > 36 and SDR > 0.54 | 3,559 | 34 |
| Drought | HI30 <= 36 and SDR <= 0.54 | 621 | 19 |
| Heat+Drought | HI30 > 36 and SDR <= 0.54 | 2,678 | 3 |

We performed GWAS independently for each phenotype under each stress condition. For each hybrid under a given stress condition, a single environment-adjusted phenotypic value was derived by fitting a fixed-effects linear model that included hybrid and nested year-location-replicate-block effects. Mean-centered hybrid coefficients were extracted as Best Linear Unbiased Estimates (BLUE) [31] and used as phenotypic inputs for GWAS. Association mapping was conducted using the multi-locus BLINK method [32], implemented in GAPIT v4.1.0 [33], with the first 5 principal components of the SNP genotype matrix included as fixed-effect covariates to account for residual population structure.

LD decay was estimated from the full unpruned G2F panel of 437,214 SNPs using pairwise squared genotype correlations (r²) computed in PLINK v1.9 [34] for SNP pairs separated by up to 300 kb. Mean r² values were calculated in 1-kb distance bins and fitted using the Hill–Weir expected-r² decay function [35,36], extended with a free background term to account for the residual long-range LD in the related hybrid panel. The resulting curve showed a background LD plateau of approximately r² = 0.16 and a half-decay distance of 15.7 kb, defined as the distance at which the fitted r² reached the midpoint between its short-range value and the background plateau (Supplementary Figure S1). Based on this estimate, we used a ±16 kb window around each prioritized SNP for downstream candidate gene annotation.

Conventional GWAS approaches often apply stringent Bonferroni or false discovery rate (FDR) corrections, which may increase false negatives and exclude biologically relevant loci [23,37]. We used a permissive threshold of p ≤ 0.05 to identify an initial set of stress-associated loci, retaining a broader pool of exploratory candidate regions for downstream sequence modeling and prioritization. Figure 1 presents an UpSet plot [38] showing the overlap among yield-associated loci across stress conditions, together with the chromosome-wise distribution of markers under each condition. Manhattan plots comparing conventional significance thresholds with the permissive threshold used in this study are shown in Supplementary Figure S2.

**Fig. 1.**
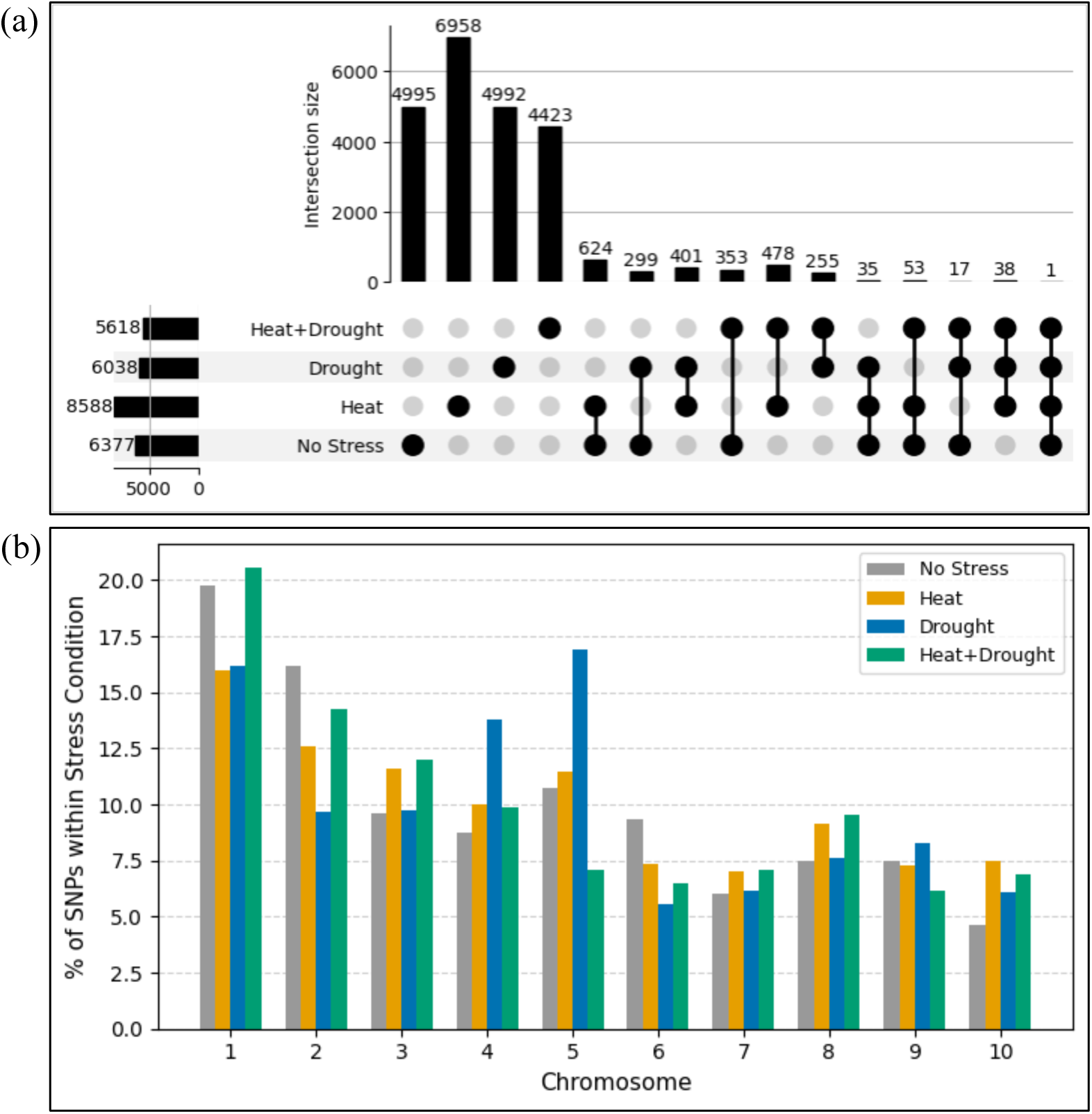
GWAS-identified candidate SNPs associated with grain yield under a permissive threshold (p ≤ 0.05). (a) UpSet plots (https://github.com/jnothman/UpSetPlot) showing the overlap of SNP sets across stress conditions; each row represents a stress condition and each column an intersection, with vertical bars indicating the number of SNPs per intersection. (b) Chromosome-wise distribution of identified loci for each stress condition.

### 2.2. Overview of GFMs

GFMs convert genomic sequences into discrete tokens that can be processed within a language modeling framework. Common tokenization strategies include single-nucleotide tokenization, in which each nucleotide is treated as an individual token [39]; fixed-length *k*-mer using overlapping or non-overlapping subsequences [40,41]; and subword-based approaches such as byte-pair encoding (BPE), which segments sequences according to frequently occurring patterns [42]. Single-nucleotide tokenization preserves base-level resolution, enabling the model to capture subtle genetic variations including SNP-level differences, whereas k-mer and BPE approaches encode short contiguous sequence patterns that may encompass local motifs and regulatory elements [14,39]. Various GFM architectures include transformer encoders (e.g., BERT [43]); transformer decoders, (e.g., GPT [44]); convolutional Hyena-based decoders [39]; and state-space models [45]. Encoder-only models are typically pre-trained using Masked Language Modeling (MLM), in which masked tokens are predicted from the surrounding sequence context [43]. On the other hand, decoder-only models use Causal Language Modeling (CLM), predicting each subsequent token autoregressively from preceding tokens [44]. Representational capacity and downstream utility of GFMs therefore depend jointly on tokenization strategy, architecture, and pre-training objective.

For instance, DNABERT [40], a BERT-based GFM pre-trained on the human genome, used overlapping *k*-mers, which capture fine-grained sequence context but introduce token redundancy, increase computational cost, and allow masked tokens to be partially inferred from adjacent unmasked k-mers. DNABERT-2 [42] addressed these limitations by replacing *k*-mers with BPE and extended pre-training on multi-species, non-plant genomes. Alternatively, Nucleotide Transformer (NT) [41] employed non-overlapping *k*-mers pre-trained on multi-species genomes, and was later adopted to plant genomes as AgroNT [17]. Other BERT-based GFMs include Grover [46], which optimized a BPE vocabulary through next-*k*-mer prediction on the human genome, and GENA-LM [47], which combined BPE with long-context mechanism based on BigBird sparse attention and recurrent memory to support inputs of up to 36 kb, with multi-species and taxon-specific variants, including *Arabidopsis*. While transformer encoders are designed to generate context-rich representations suitable for classification and regression tasks, decoder-based GFMs target autoregressive sequence generation. MegaDNA [48] applied multi-scale transformers to bacteriophage genomes at single-nucleotide resolution, and DNAGPT [49] was pre-trained on mammalian genomes using *k*-mer tokenization. Genos [50] integrated a transformer decoder with a mixture-of-experts architecture to model human genomes at single-nucleotide resolution over contexts of up to 1 Mb.

Beyond transformer-based models, GPN [51] used a convolutional neural network with one-hot encoded inputs (each nucleotide represented as a binary vector) for genome-wide variant effect prediction across multiple species. HyenaDNA [39], pre-trained on the human genome, employed the convolutional Hyena architecture to support context lengths of up to 1 Mb at single-nucleotide resolution. Evo 2 [52] extended this approach by using multi-hybrid StripedHyena-2 architecture, and was pre-trained on genomic sequences spanning all domains of life. Caduceus [45], pre-trained on the human genome at nucleotide resolution, extended the Mamba state-space architecture through bidirectional modeling and reverse-complement equivariance. Building on this, PlantCaduceus [53] was pre-trained on angiosperm genomes and demonstrated cross-species transferability to maize after fine-tuning on a small labeled *Arabidopsis* dataset across several downstream tasks.

### 2.3. Environmental Context Adaptation of GFM

After reviewing existing GFMs, we selected AgroNT as the backbone model because its pre-training corpus spans 48 plant species, including maize, making it well-suited for modeling plant regulatory sequence context and adapting genomic representations for maize stress-response analysis. The overall workflow is illustrated in Fig. 2.

**Fig. 2.**
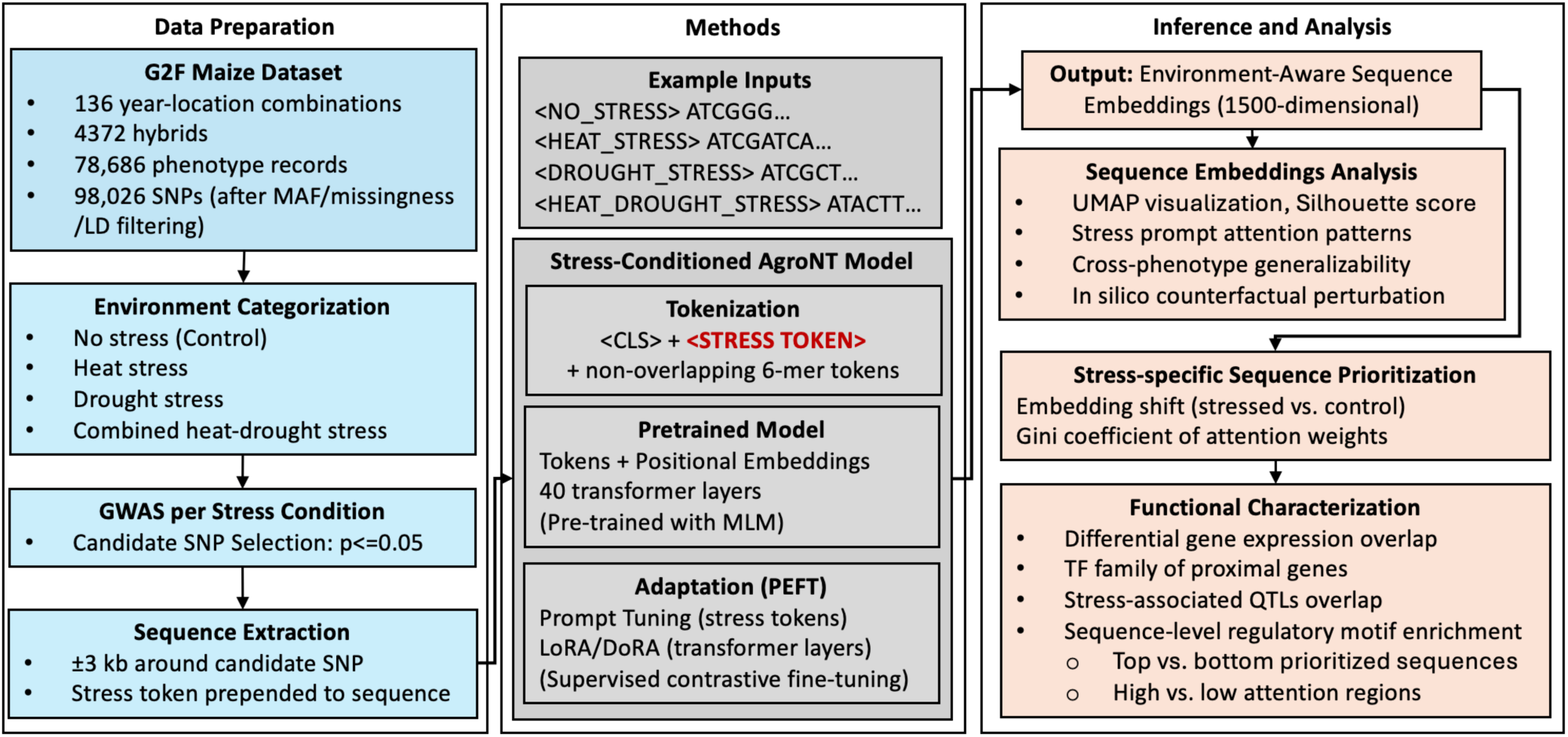
Environment-aware DNA-LLM framework for prioritizing stress-responsive genomic regions in maize. Genotype-phenotype data were stratified by environmental stress condition, followed by stress-specific GWAS, stress-conditioned AgroNT adaptation, embedding-based sequence prioritization, and downstream functional characterization.

#### 2.3.1. Pre-trained AgroNT

AgroNT [17] is a BERT-based encoder framework that adopts a non-overlapping 6-mer tokenization strategy with a vocabulary of 4,105 tokens. It accepts sequences of up to 6,144 bp, corresponding to 1,024 DNA tokens, which are mapped to 1,500-dimensional embeddings, combined with their positional embeddings, and subsequently processed through 40 transformer layers to generate contextual representations. Each layer applies multi-head self-attention, in which attention weights reflect how strongly each token attends to other tokens. The pre-training was performed using an MLM objective on approximately 10.5 million DNA sequences from the Ensembl Plants database [54].

#### 2.3.2. Stress-Conditioned AgroNT

We introduced four stress-specific prompt tokens, <NO_STRESS>, <HEAT_STRESS>, <DROUGHT_STRESS>, and <HEAT_DROUGHT_STRESS>, into the AgroNT vocabulary as special tokens. Sequences spanning ±3 kb around each GWAS-identified locus were extracted for all stress conditions to remain within AgroNT’s maximum input length, yielding approximately 1,000 DNA tokens per sequence. Each genomic sequence was prepended with its corresponding stress prompt to condition the model on the environmental context. During tokenization, AgroNT automatically inserted a <CLS> token at the beginning of each input, with the stress token positioned immediately after it.

Environmental context could alternatively be incorporated through continuous prompt-based conditioning, such as soft prompts or prefix tuning [55,56], or through condition-specific adapter modules [57]. In our formulation, explicit vocabulary tokens provide a discrete, learned representation of environmental context that modulates DNA-token representations throughout the transformer layers. We adopted PEFT for stress-conditioned adaptation of AgroNT.

##### PEFT

These strategies update only a small subset of model parameters while keeping the pre-trained backbone frozen. We applied a prompt tuning (PT) approach [55], optimizing only the 1,500-dimensional *input embedding vectors* of the stress tokens, resulting in 4×1,500 trainable parameters. To further adapt the internal sequence representations, we combined PT with low-rank adaptation modules inserted into the *transformer layers*. We evaluated LoRA [58], which parameterizes weight updates using trainable low-rank matrices, and DoRA [59], which decomposes pre-trained weights into magnitude and directional components to improve learning capacity and training stability compared with LoRA.

##### Fine-tuning Method

Fine-tuning was conducted using yield-associated loci, while loci on chromosomes 8 and 9 were excluded from training and retained as an independent test set to evaluate generalization across genomic regions. Training was performed for 10 epochs using the AdamW optimizer [60] with the learning rate of 0.0001 for the stress token embeddings and a global batch size of 32 sequences (approximately 32K DNA tokens per optimization step). For LoRA and DoRA adaptation, the low-rank dimension was set to *r*=8, with a scaling factor α=16 and dropout of 0.05. The low-rank modules were applied to the query and value projections of the transformer layers, while the remaining pre-trained AgroNT parameters were kept frozen. Additionally, a 3-epoch warm-up strategy was used, during which only the stress-token embeddings were optimized. The LoRA/DoRA parameters were subsequently enabled for the remaining 7 epochs using a learning rate of 0.00005, while the stress-token embeddings continued to be optimized at 0.0001.

##### Fine-tuning Objective

We applied the supervised contrastive loss [61] to the *final transformer-layer embedding of the stress token*, which served as a stress-conditioned summary representation of each sequence. This objective encouraged representations from the same stress condition to lie closer in the embedding space than those corresponding to different stress conditions. The language-modeling head used during MLM pre-training was not used during fine-tuning.

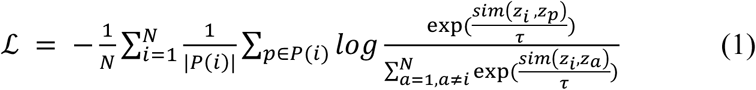

where, *z_i_*, *z_p_*, *z_a_* denote normalized final-layer stress token embeddings of the anchor sequence *i*, a positive sequence *p*, and comparison sequence *a*, respectively. The anchor is the reference sample for which the loss is computed. Positive sequences are other samples within the same mini-batch that share the anchor’s stress condition, whereas comparison sequences include all samples in the mini-batch except the anchor. *P*(*i*) denotes the set of positive samples for anchor *i*; N denotes the mini-batch size; *sim*(⋅,⋅) denotes cosine similarity; and *τ* is the temperature parameter, set to 0.07.

### 2.4. Stress-specific Sequence Prioritization

Next, we formulated a prioritization strategy to rank sequences independently under heat, drought, and combined heat-drought stress. For each sequence, the *final transformer-layer embeddings of the 6-mer DNA tokens* were extracted from the stress-conditioned AgroNT model, excluding the <CLS> and stress tokens. Sequence-level representations were obtained by mean-pooling the DNA-token embeddings, resulting in a 1,500-dimensional vector per sequence. A priority score was then computed by integrating two components.

a. **Embedding shift score (stress vs. control):** The embedding shift was calculated as the cosine distance between sequence representations generated using the corresponding stress prompt and the <NO_STRESS> prompt:

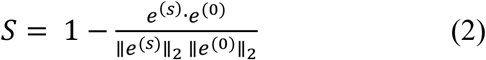

where, *e*^(*s*)^ and *e*^(0)^ denote the mean-pooled DNA-token embeddings of the same genomic sequence under the corresponding stress and no-stress conditions, respectively. Larger values indicate greater stress-dependent changes in the sequence representation.
b. **Gini coefficient of attention weights:** The Gini coefficient was calculated from the final-layer attention weights assigned by the stress token to DNA tokens, averaged across the 20 self-attention heads. After excluding the <CLS> and stress-token positions, the attention weights over the remaining DNA tokens were normalized to sum to one, and the Gini coefficient was calculated as:

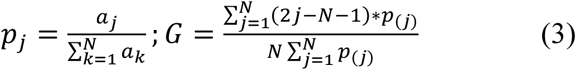

where, *a_j_* is the normalized attention weight assigned to the *j^t^*^ℎ^ DNA token, *p_j_* is the normalized attention weight, N is the number of DNA tokens, and *p_(j)_* denotes the normalized attention weights sorted in the ascending order. Higher Gini coefficients indicate that attention was concentrated on a smaller subset of DNA tokens rather than distributed uniformly across the sequence. Because regulatory signals may be localized within specific sequence segments, this measure was used to identify sequences containing focal regions emphasized by the model.

The priority score, ranging from 0 to 1, was calculated as the geometric mean of the percentile ranks of the above two components:

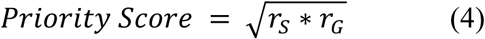

where, *r_S_* and *r_G_* denote the percentile ranks of the embedding shift score and attention Gini scores, respectively. Therefore, higher priority scores indicate sequences exhibiting both larger stress-induced representation shifts and more concentrated stress-token attention patterns.

## 3. Results

### 3.1. Environment-Aware Genomic Representations

We first evaluated whether stress-prompt conditioning reshaped AgroNT’s sequence representations (mean-pooled 6-mer DNA token embeddings from the final transformer layer) relative to the pre-trained model by examining stress-dependent clustering patterns, generalizability across phenotypes, and the modulations in sequence representations across environmental contexts.

#### 3.1.1. Clustering Patterns of Sequence Embeddings

##### UMAP

We projected the 1,500-dimensional sequence embeddings into two dimensions using Uniform Manifold Approximation and Projection (UMAP) [62] to qualitatively assess stress-associated clustering patterns (Fig. 3). Embeddings from the pre-trained model largely overlapped across stress conditions, indicating little stress-specific structure. Following PT, the embeddings formed distinct clusters by stress condition, showing that AgroNT learned stress-conditioned sequence representations through prompt-based adaptation. Comparable clustering patterns in the training and held-out test chromosomes suggest that this stress-conditioned structure generalized beyond the chromosomes used for fine-tuning rather than reflecting chromosome-specific patterns. However, because PT already produced clear separation in the two-dimensional projection, UMAP visualization alone was insufficient to determine whether LoRA or DoRA augmentation further improved the separation of stress-conditioned representations.

**Fig. 3.**
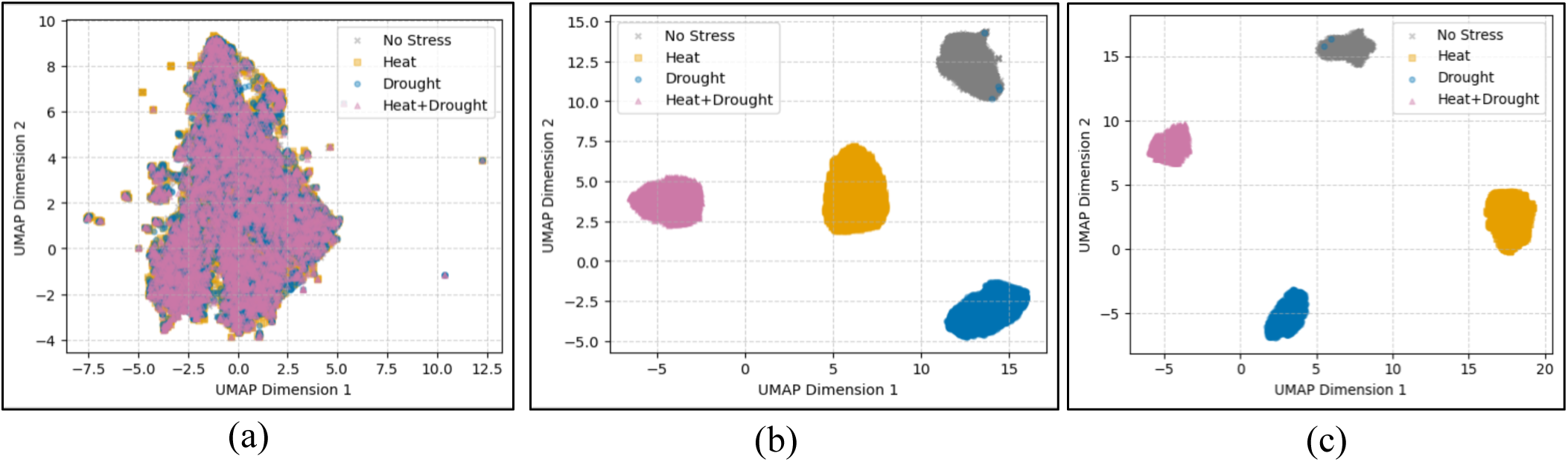
UMAP of yield-associated sequence embeddings from the pre-trained and stress-conditioned AgroNT models. Each sequence is represented by mean-pooled final-layer embeddings of its 6-mer DNA tokens. (a) Pre-trained model embeddings without stress prompt conditioning, showing no discernible separation across stress conditions. (b) Stress-conditioned embeddings after PT on the training set (chromosomes 1–7 and 10), showing stress-associated clustering. (c) Stress-conditioned embeddings on the held-out test set (chromosomes 8 and 9), showing a similar clustering pattern across stress conditions and suggesting that the learned structure was not driven by chromosome-specific overfitting.

##### Silhouette Score

Clustering quality was quantified using the silhouette score [63], with stress conditions as class labels and cosine distance computed in the 1,500-dimensional sequence-embedding space (Table 2). The score compares within-condition cohesion with separation from other conditions, ranging from −1 (samples closer to a different cluster) to 1 (well-separated clusters). The near-zero score derived from the pre-trained model embeddings indicated substantial overlap across stress conditions, whereas PT significantly improved cluster separation. Augmenting PT with LoRA or DoRA produced further improvements, with PT+DoRA resulting in the strongest separation.

**Table 2.** Silhouette score evaluation of yield-associated sequence embeddings from the pre-trained and stress-conditioned AgroNT models. Scores approaching +1 indicate well-separated stress-condition clusters, whereas scores near 0 indicate substantial overlap.

| **Split Set** | **Pre-trained AgroNT<br>(without stress prompts)** | **Stress-Conditioned AgroNT** |  |  |
|---|---|---|---|---|
|  |  | **PT** | **PT+LoRA** | **PT+DoRA** |
| Training | -0.02 | 0.40 | 0.47 | 0.62 |
| Test |  | 0.41 | 0.48 | 0.63 |

#### 3.1.2. Cross-Phenotype Generalizability

To assess whether the learned stress-conditioned representations transferred beyond grain yield, the model fine-tuned on yield-associated loci was applied to flowering-trait-associated loci without additional model updates. As observed for the yield loci, pre-trained sequence embeddings of ±3kb regions surrounding flowering-trait loci overlapped substantially across stress conditions, whereas the stress-conditioned model produced distinct stress-specific clusters (Supplementary Fig. S3). Silhouette scores were approximately 0.41 (PT), 0.47 (PT+LoRA), and 0.62 (PT+DoRA) across all flowering traits, indicating that the learned stress-conditioned embedding structure generalized to phenotypes not used during fine-tuning.

#### 3.1.3. Environmental Context Modulation Analysis

Next, we examined how stress prompts modulated the representations and attention patterns of the same genomic sequences, providing the basis for the two model-derived signals integrated into the priority score.

##### In Silico Counterfactual Context Perturbation

We first evaluated whether the learned sequence representations were modulated by the stress prompt when the underlying genomic sequence was held fixed, particularly because many loci were uniquely associated with individual stress conditions rather than shared across environments (Fig. 2). For each sequence under heat, drought, and heat-drought stress, we replaced the corresponding stress prompt with <NO_STRESS> while keeping the nucleotide sequence unchanged. Following prompt substitution, these representations shifted toward the <NO_STRESS> representation cluster (Supplementary Fig. S4), indicating that the stress prompt influenced the learned sequence representations. Importantly, the magnitude of this shift, measured by cosine distance, varied across sequences, suggesting that the effect of environmental conditioning depended on sequence content rather than reflecting a uniform, sequence-independent contribution of the stress prompt. These sequence-specific representation shifts were subsequently used in the priority score.

##### Stress Prompt Attention Analysis

Secondly, we evaluated how attention patterns varied across stress prompts for the same genomic sequence using SNP S2_33499321, a GWAS locus shared across all four stress conditions (Fig. 2). We observed that the final-layer attention weights from the stress token to the DNA tokens differed across stress contexts (Fig. 4), indicating prompt-dependent redistribution of attention within the same nucleotide sequence. Hence, we quantified the concentration of stress-token attention using the Gini coefficient, with higher values indicating that attention was concentrated on a smaller subset of sequence regions. This attention concentration measure was also incorporated in the priority score.

**Fig. 4.**
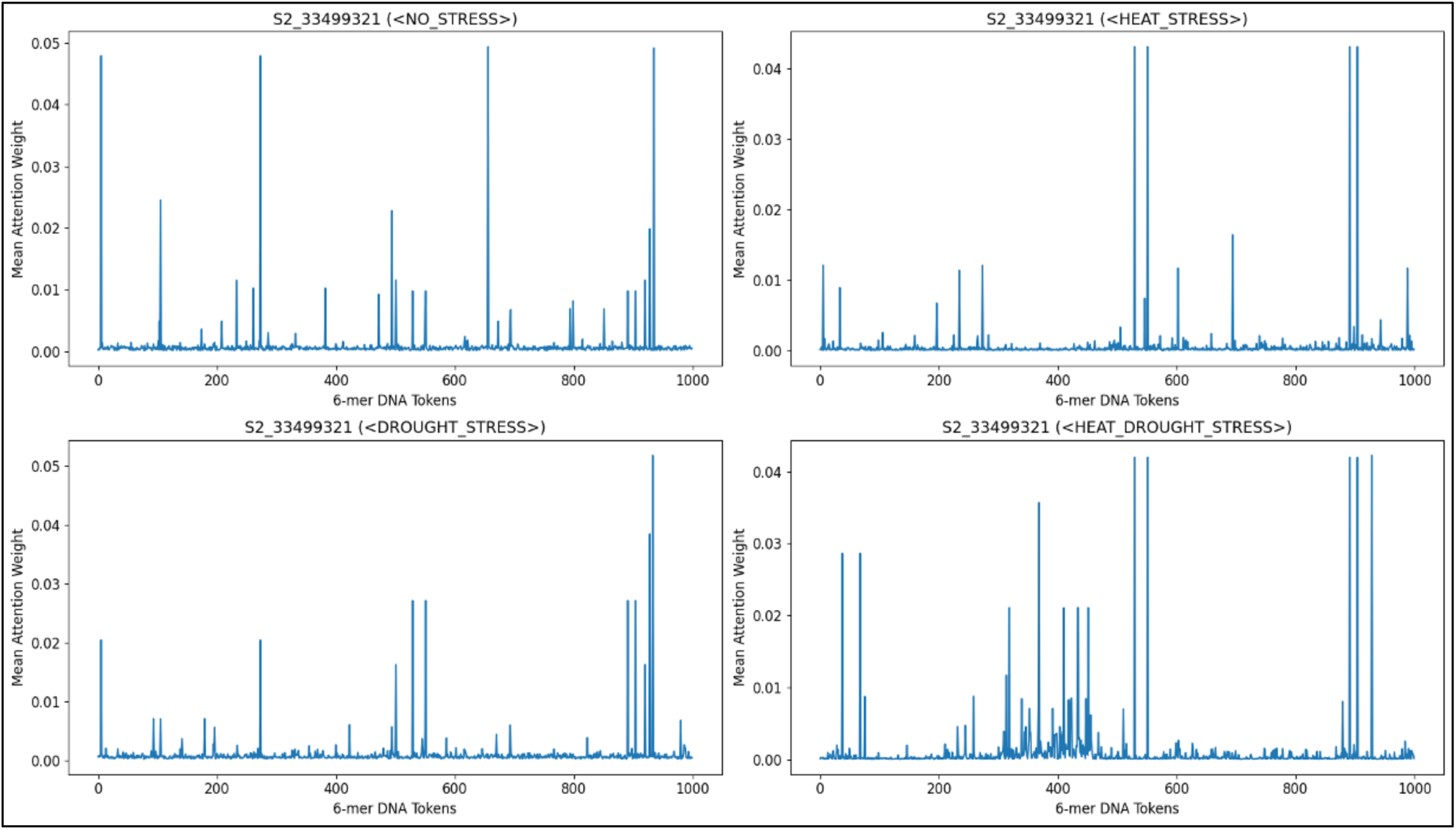
Stress-prompt attention across 6-mer DNA tokens for a shared GWAS locus. Mean attention weights, averaged across 20 heads in the final transformer layer, from each stress token to 6-mer DNA tokens are shown for marker S2_33499321, the only grain yield-associated locus identified at p ≤ 0.05 under all four stress conditions. Differences in attention patterns across prompts illustrate stress-dependent contextual modulation of the same genomic sequence.

### 3.2. Stress-Specific Sequence Prioritization

For yield-associated sequences under heat, drought, and heat-drought conditions, priority scores were computed using Equation (4), and *the top 10% were selected as prioritized candidates* for downstream functional characterization. Since PT+DoRA achieved a higher silhouette score than PT+LoRA (0.62 vs. 0.47; Table 2), it was selected for subsequent analyses, while PT was retained for comparison.

We generated Manhattan-style plots of the priority scores across GWAS loci (p ≤ 0.05), highlighting overlap between the top 10% prioritized candidates and the top 10% GWAS-ranked loci with the smallest p-values (Fig. 5). Priority scores showed limited correspondence with GWAS-based rankings, suggesting that the model-based prioritization captured information not fully reflected by conventional statistical association analysis. The top 10% of GWAS-ranked loci were therefore included as a reference set for gene-level and MQTL-overlap analyses.

**Fig. 5.**
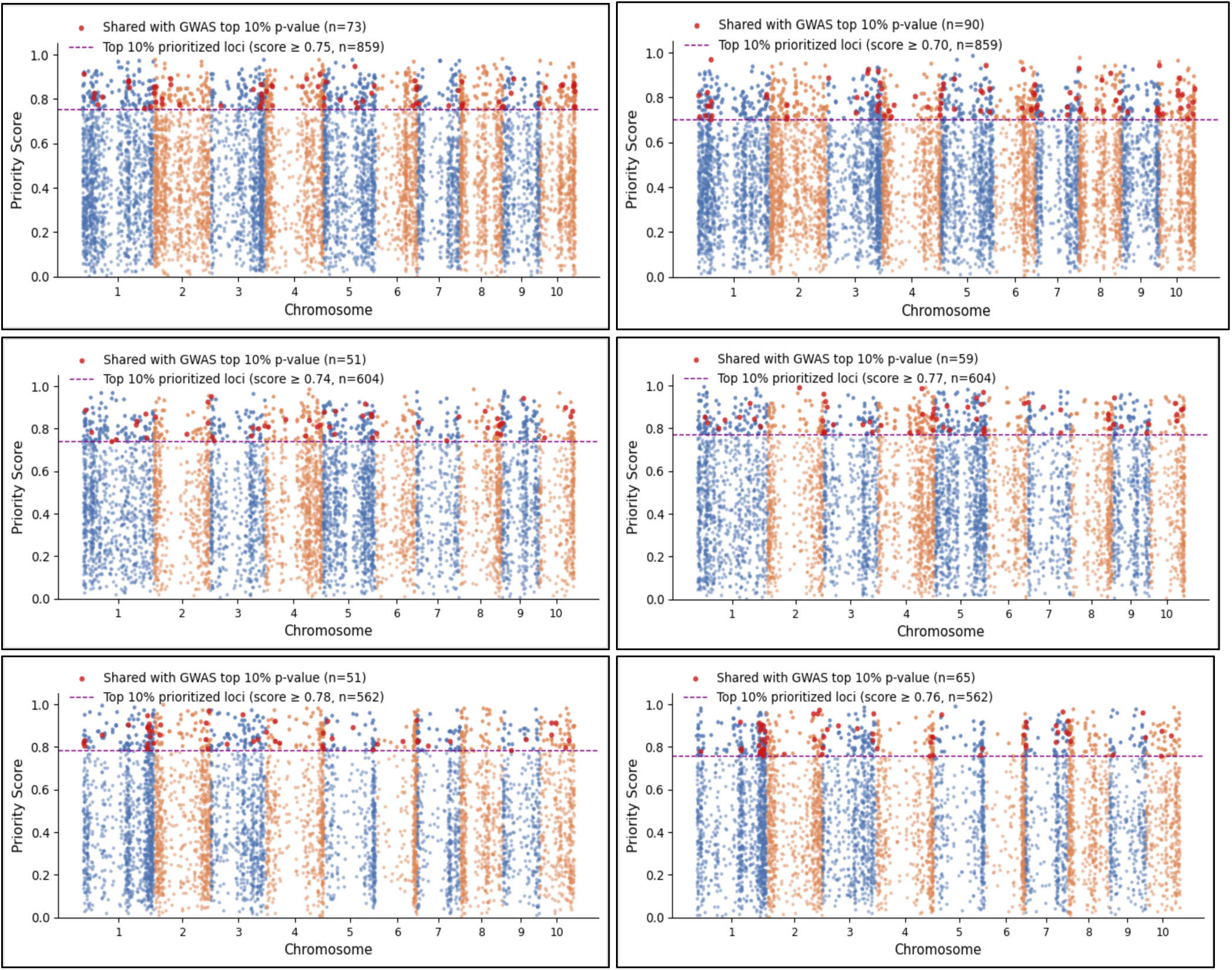
Manhattan-style plots of stress-conditioned AgroNT-based priority scores. Rows correspond to heat, drought, and combined heat-drought stress (top to bottom); columns correspond to PT and PT+DoRA (left to right). The dashed line marks the top 10% priority-score threshold. Red points indicate loci shared between the top 10% model-prioritized candidates and the top 10% GWAS-ranked loci by p-value.

### 3.3. Functional Characterization of Prioritized Sequences

#### 3.3.1. SNP Annotation

SNPs were annotated using SnpEff v5.4a [64], with proximal genes defined within the LD half-decay distance (∼16 kb, Fig. S1). Prioritized loci spanned coding and noncoding regions, with exonic, upstream, and downstream annotations frequently represented alongside intergenic loci (Fig. 6a–c). Proximal genes near prioritized loci showed limited overlap across stress conditions (Fig. 6d–e), supporting stress-dependent prioritization patterns. Notably, the PT and PT+DoRA proximal-gene sets showed low Jaccard similarity (intersection over union; ∼9%; Fig. 6f), indicating that DoRA augmentation prioritized largely distinct candidates. The functional relevance of these candidates is assessed in the following analyses.

**Fig. 6.**
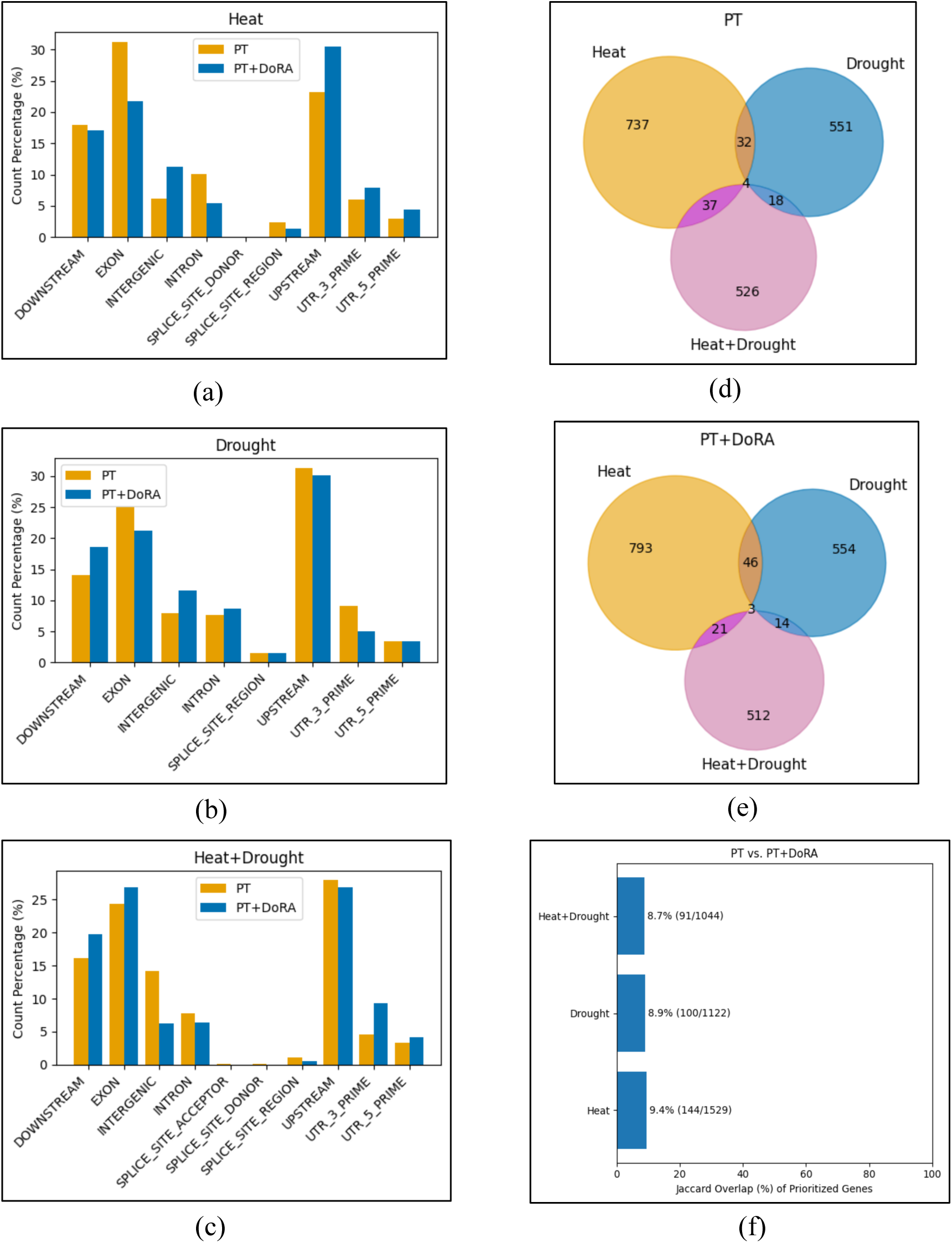
Genomic annotation and comparative analysis of prioritized loci. (a–c) Genomic feature distribution of the top 10% prioritized loci under heat, drought, and combined heat-drought stress, compared between PT and PT+DoRA. Coding-sequence effects reported by SnpEff (missense, synonymous, start/stop gain or loss, and related variants) were collapsed into a single “exon” category; other effects retained their reported genomic feature. (d–e) Overlap of proximal genes across stress conditions for each method. (f) Jaccard similarity (%) between PT and PT+DoRA proximal-gene sets across stress conditions; low values indicate that the methods prioritize largely distinct gene sets.

#### 3.3.2. Transcriptomic Overlap Analysis

We first analyzed whether genes proximal to prioritized loci under heat and drought stress overlapped spatiotemporal stress-responsive differentially expressed genes (DEGs) curated from published maize transcriptomic studies. Gene identifiers from different sources were harmonized using MaizeMine v1.6 [65].^3^ When multiple prioritized proximal genes mapped to the same DEG, the DEG was counted only once to avoid inflation from many-to-one mappings.

##### Heat-Prioritized Proximal Genes

Genes proximal to heat-prioritized loci were compared against transcriptomic data from B73 leaf, tassel, ear, and silk tissues collected at 2 and 48 hours after heat stress (HAH) during the reproductive (R1) stage [66]. PT+DoRA showed a higher proportion of DEGs than PT on average across tissue-duration combinations, and both exceeded GWAS in several conditions (Fig. 7a). The low Jaccard similarity (6–13%, Fig. 7b) between PT and PT+DoRA DEG sets indicated that the two strategies captured largely complementary subsets of heat-responsive genes. *PT+DoRA also identified DEGs with greater mean absolute expression changes (|log2FC|) than PT in most conditions (Fig. 7c), suggesting that its prioritized loci were associated with a subset of genes showing stronger transcriptional responses.* Overlapping DEGs included both positive and negative log2-fold changes (Supplementary Fig. S5a-b), while hierarchical clustering separated them into modules with distinct spatiotemporal profiles (Supplementary File 2), linking prioritized loci to diverse early and late heat-response programs rather than a single uniform expression pattern (Supplementary Fig. S5c).

**Fig. 7.**
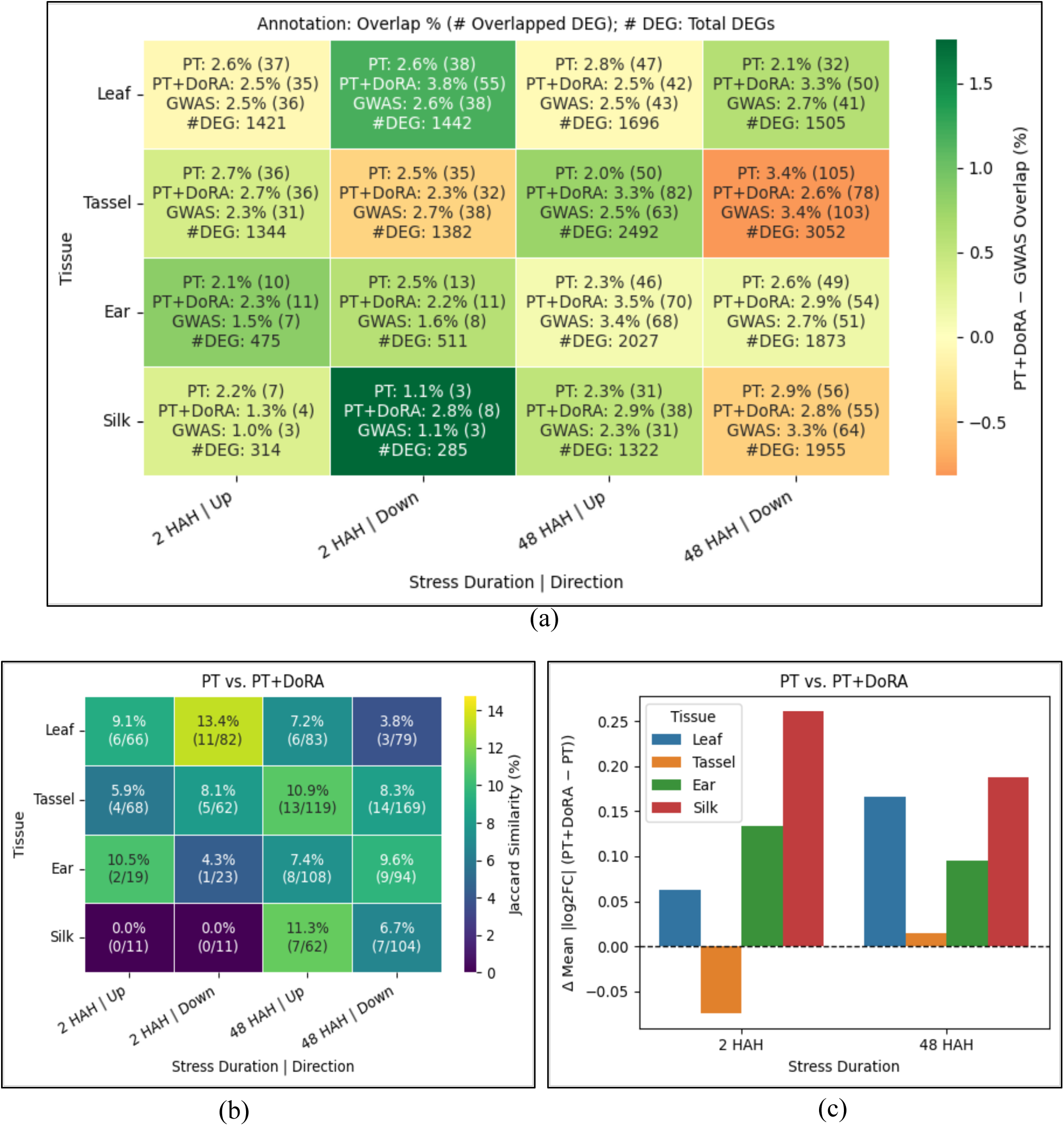
Comparative analysis of heat-prioritized proximal genes with heat-stress transcriptomic data. (a) Overlap (%) of genes proximal to the top 10% loci with heat-responsive DEGs, shown for PT, PT+DoRA, and GWAS ranking across tissues and stress durations (HAH, hours after heat); cell color reflects the PT+DoRA − GWAS overlap difference, with positive values indicating higher overlap for PT+DoRA. (b) Jaccard similarity (%) between the DEG sets recovered by PT-and PT+DoRA-prioritized loci. (c) Difference in mean |log₂FC| between DEGs recovered by PT+DoRA-and PT-prioritized loci; positive values indicate that DoRA augmentation recovers DEGs with larger expression changes.

During the tetrad stage of pollen development of B73 genotype, 169 up-regulated and 131 down-regulated DEGs were reported [67]; PT, PT+DoRA, and GWAS overlapped with 4/1, 3/3, and 2/5 up/down-regulated genes, respectively. In another study of kernel development under heat stress, 338 up-regulated and 432 down-regulated DEGs were shared between heat-tolerant (PF5411-1) and heat-sensitive (LH150) genotypes [68]; PT, PT+DoRA, and GWAS overlapped with 7/16, 4/15, and 6/8 of these genes, respectively. Although these genotypes were not represented in the G2F population, the observed overlaps provide additional evidence that some prioritized loci are associated with heat-responsive genes identified across distinct genetic backgrounds.

##### Drought-Prioritized Proximal Genes

Comparison with published drought transcriptomic datasets [69–71] showed that PT+DoRA recovered more DEGs than PT and GWAS in B73 leaf, tassel and ear tissues during reproductive stage (Table 3). Further, the union of genes recovered by PT and PT+DoRA consistently captured a broader set of drought-responsive DEGs than either method alone across studies, further supporting their complementary nature.

**Table 3.** Overlap of genes proximal to yield-associated prioritized loci under drought stress with published DEGs.

| **Genotype/<br>Growth<br>Stage** | **Tissue** | **Regulation** | **# DEGs** | **# Overlap of Top 10%<br>DNA-LLM Candidates** |  |  | **# Overlap of Top<br>10% GWAS<br>Candidates** |
|---|---|---|---|---|---|---|---|
|  |  |  |  | **PT** | **PT + DoRA** | **Union** |  |
| B73/R1[69] | Leaf | - | 601 | 6 | 10 | 13 | 8 |
|  | Tassel | - | 9377 | 117 | 120 | 215 | 107 |
|  | Ear | - | 2595 | 35 | 39 | 69 | 32 |
| B73/R1 [71] | Ovary | Up | 57 | 2 | 2 | 4 | 3 |
|  |  | Down | 74 | 2 | 1 | 3 | 4 |
| B73/VT [70] | Leaf | Up | 284 | 2 | 2 | 4 | 2 |
|  |  | Down | 229 | 5 | 5 | 10 | 6 |

#### 3.3.3. TF Family of Proximal Genes

Maize TF annotations were obtained from Grassius^4^ [72] to evaluate whether proximal genes near heat-, drought-, and heat-drought-prioritized loci belonged to TF families implicated in abiotic stress responses. After excluding unclassified (“Orphans”) TF genes, PT+DoRA identified 190 TF-encoding genes across the three conditions, compared with 159 for PT and 163 for the GWAS reference set. These genes spanned several stress-associated TF families, including **bHLH, Homeobox, MYB, bZIP, NAC, WRKY, AP2/ERF, and MADS-box** [68,73–76]. Among the three approaches, PT+DoRA identified the largest number of TF genes under heat (81, compared with 59 for PT and 56 for GWAS) and drought (63, compared with 58 and 55, respectively), whereas the GWAS reference set identified the largest number under heat-drought stress (52, compared with 42 for PT and 46 for PT+DoRA). Overall, PT+DoRA resulted in a larger number TF-encoding genes associated with stress-related TF families (Fig. 8).

**Fig. 8.**
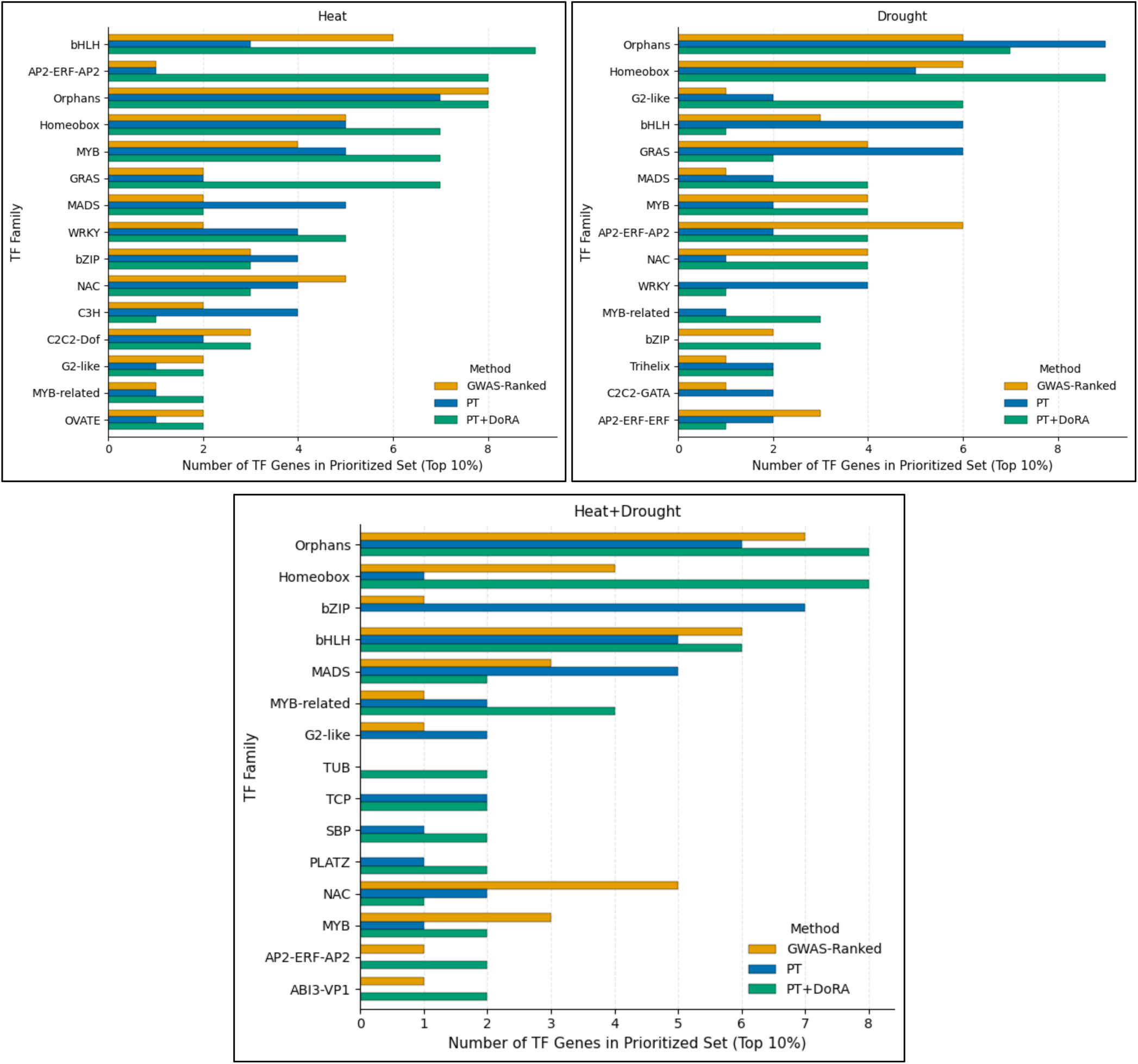
TF family composition of genes proximal to prioritized loci under heat, drought, and combined heat-drought stress. Results are shown for PT, PT+DoRA, and GWAS ranking. TF families are ranked by the maximum number of genes recovered across the two DNA-LLM-based methods and top 15 families are shown.

#### 3.3.4. Overlap and Enrichment with Stress-Associated MQTLs

To assess whether prioritized loci overlapped previously reported abiotic stress-associated regions, we compared them with published maize MQTLs [77], which integrate QTL evidence across studies, populations, and environments. MQTL coordinates were lifted from B73 v4 to v5 using the MaizeGDB^5^ chain file and CrossMap v0.7.0 [78] and BEDTools [79]. Because lift-over fragmented some large MQTL intervals, fragments from each MQTL were consolidated into a single v5 interval spanning the outer bounds of the dominant contiguous mapping cluster.

Under heat stress, PT recovered one SNP (S1_20257139) within one of 5 heat-associated MQTLs, whereas PT+DoRA recovered 3 SNPs (S1_14258495, S2_14209026, S9_112798223), each overlapping a different heat-associated MQTL. Under drought stress, PT recovered 34 prioritized SNPs within 13 of 22 drought-associated MQTLs, and PT+DoRA 27 SNPs within 8 MQTLs. Under heat-drought stress, PT recovered 23 SNPs within 11 of 32 abiotic-stress-associated MQTLs (1 heat-and 9 drought-associated), and PT+DoRA 18 SNPs within 16 MQTLs (2 heat- and 11 drought-associated). The overlap of combined-stress-prioritized loci with MQTLs associated with both individual stresses is consistent with partially shared genomic responses.

As drought-associated MQTLs constituted the largest stress-specific set and showed substantial overlap with prioritized loci, we performed a permutation-based enrichment analysis for drought (10,000 iterations). Size-matched random SNP sets were sampled without replacement from the GWAS candidate background (p ≤ 0.05), and an empirical one-sided p-value was calculated as (r + 1)/(n + 1), where r is the number of iterations with MQTL overlap at least as large as that observed. *PT showed significant enrichment, with 34 observed overlapping SNPs compared with 20.8 expected under random sampling (1.63-fold enrichment, empirical p = 0.0042). PT+DoRA showed a 1.30-fold enrichment that was not statistically significant (p = 0.108).* For comparison, a size-matched set of top GWAS-ranked loci showed significant but weaker enrichment, with 31 overlapping SNPs (1.49-fold enrichment, p = 0.0218).

#### 3.3.5. Regulatory Motif Enrichment Analysis

To further interpret the regulatory relevance of DNA-LLM-based prioritization, we conducted Analysis of Motif Enrichment (AME; MEME Suite (v5.5.5) [80] using the JASPAR 2026 plant motif database [81], both at the sequence level (full ±3 kb windows) and within high-attention regions of the prioritized sequences. The latter analysis provided additional insight into the sequence patterns emphasized by the model. In addition to PT and PT+DoRA, PT+LoRA results were included to compare adaptation-specific motif profiles. We also evaluated score ablations based on the individual components of the priority score using PT+DoRA (Equation 2 and 3).

##### Motif Enrichment in Prioritized Sequences

The top 10% prioritized sequences were compared with the bottom 10% low-priority sequences using the full ±3 kb DNA-LLM input windows to identify TF motif families preferentially represented among prioritized sequences. *As foreground and background sets were defined by the priority score and were not explicitly matched for GC content, these results were interpreted as method-specific motif profiles rather than GC-controlled enrichment estimates.* AME identified significantly enriched motifs (FDR ≤ 0.05) corresponding to TF families with reported roles in abiotic stress responses, spanning **ERF/DREB, bHLH, NAC, MYB, WRKY, bZIP, ARF, HD-ZIP, DOF, MADS-box type II,** and **CAMTA** [74,76,82–87] (Fig. 9). Enrichment profiles varied across adaptation methods and prioritization strategies.

**Fig. 9.**
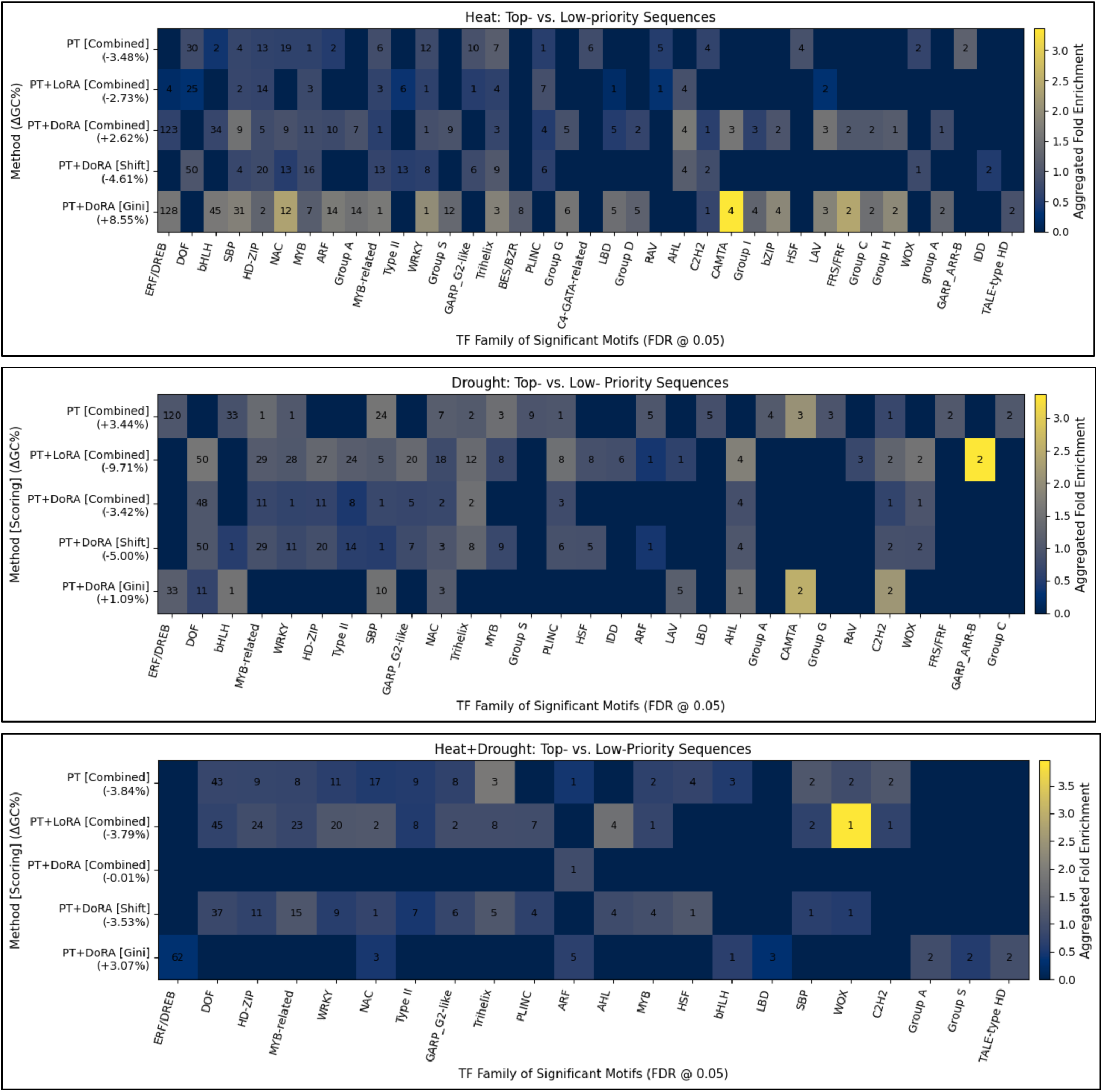
TF families of significant motifs enriched in prioritized sequences relative to low-priority sequences under heat, drought, and combined heat-drought stress. Results are shown for PT, PT+LoRA, and PT+DoRA. “Combined” denotes the prioritization score defined in Equation 4, whereas “Shift” and “Gini” denote ablation analyses using the individual score components (Equation 2 and 3). Only TF families containing at least one significant motif in any method are shown. Color indicates the log₂-transformed aggregated fold enrichment for each TF family, calculated as the ratio of foreground to background sequence matches summed across all significant motifs in that family; ΔGC values in the row labels give the foreground−background GC difference.

**Fig. 10.**
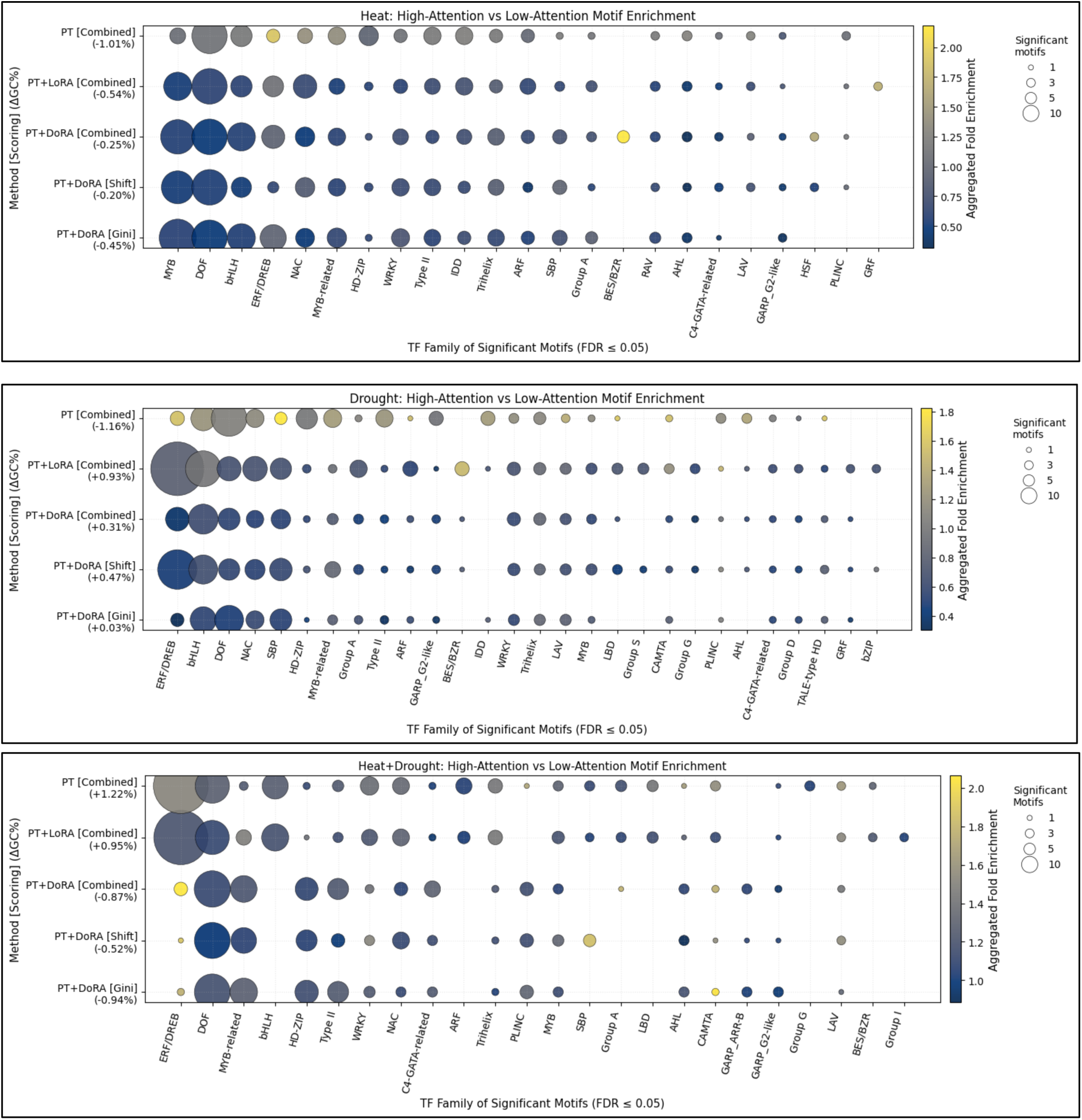
Significant TF motif families enriched within high-attention regions of prioritized sequences, relative to GC-matched low-attention regions from the same sequences, under heat, drought, and combined heat–drought stress. Results are shown for PT, PT+LoRA, and PT+DoRA. “Combined” denotes the prioritization score defined in Equation 4, whereas “Shift” and “Gini” denote ablation analyses using the individual score components (Equation 2 and 3). Only TF families containing at least three significant motifs in any method are shown. Bubble size indicates the number of significant motifs within each TF family. Color indicates the log₂-transformed aggregated fold enrichment for each TF family, calculated as the ratio of total foreground to total background sequence matches across all significant motifs in that family; ΔGC values in the row labels give the foreground−background GC difference.

Under heat stress (Fig. 9a), PT primarily recovered motifs associated with DOF, NAC, HD-ZIP, and WRKY families, whereas PT+DoRA favored ERF/DREB, bHLH, NAC, MYB, ARF, and CAMTA motifs. PT+LoRA predominantly recovered DOF, HD-ZIP, and MADS-box type II motifs. All three adaptation strategies showed modest GC imbalance. Score ablation analysis indicated that embedding-shift only prioritization favored AT-rich sequences (ΔGC = −4.61%), whereas Gini-based prioritization favored GC-rich sequences (ΔGC = +8.55%) and recovered a broader set of motifs. Because several motifs identified under Gini-based prioritization, including ERF/DREB and bHLH, have GC-rich consensus patterns, part of the observed enrichment may reflect sequence-compositional bias rather than stress-associated signal alone.

Under drought stress (Fig. 9b), PT predominantly recovered ERF/DREB, bHLH, and SBP motifs, whereas PT+DoRA recovered DOF, HD-ZIP, and MYB-related motifs, with both showing modest GC imbalance. PT+LoRA recovered the broadest motif profile, including DOF, MYB-related, WRKY, MADS-box type II, HD-ZIP, and Trihelix; however, this broader recovery coincided with a substantial GC imbalance (ΔGC = −9.71%). Score ablation analysis showed that the combined priority score reduced the compositional imbalance observed for embedding-shift prioritization (ΔGC = −5.00%), while Gini-based prioritization was comparatively well balanced (ΔGC = +1.09%).

Under heat-drought stress (Fig. 9c), PT and PT+LoRA recovered broad and overlapping motif profiles, including DOF, NAC, WRKY, MYB-related, and HD-ZIP motifs, whereas PT+DoRA recovered only an ARF motif. The combined score achieved the closest GC balance (ΔGC = −0.01%), compared with embedding shift (−3.53%) and Gini-based prioritization (+3.07%), suggesting that the corresponding motif profile was less likely to be strongly influenced by GC-compositional differences. Unlike heat stress, but consistent with drought, embedding-shift prioritization recovered more motif families than Gini-based prioritization.

##### Regulatory Motifs Enriched in High-Attention Regions

For each prioritized sequence, DNA tokens receiving attention scores in the top 5% from the stress token were designated as high-attention tokens. Their genomic intervals were extended by ±30 bp and merged when overlapping or adjacent to generate non-redundant foreground regions. Background regions were selected from the bottom 20% of attention scores within the same sequences, excluding regions that overlapped the foreground, and then matched to the foreground by GC content. *AME was used to test whether these model-emphasized regions were enriched (FDR ≤ 0.05) for known plant regulatory motifs, including motifs associated with stress-responsive TF families*.

Using this attention-guided analysis, high-attention regions across all stress conditions showed broad enrichment for TF families associated with abiotic-stress responses, including **ERF/DREB, bHLH, DOF, NAC, MYB-related, HD-ZIP,** and **WRKY** (Fig. 8) [74,82–84]. *Heat-emphasized regions additionally showed strong recovery of MYB motifs, and drought-emphasized regions showed enrichment of SBP motifs.* Differences among PT, PT+LoRA, and PT+DoRA further indicated that the adaptation strategy influenced the regulatory features emphasized by the model. Importantly, because foreground and background regions were closely GC-matched (|ΔGC| ≤ 1.22% across all methods and conditions), these enrichments were unlikely to be driven by sequence-compositional differences than the sequence-level comparisons described above.

#### 3.3.6. Summary of Prioritized Loci

As the downstream analyses supported the biological relevance of both PT and PT+DoRA, Supplementary File 1 presents *the union of loci within the top 10% by either method* across all stress conditions. Each locus was annotated with its genomic feature (Section 3.3.1), transcriptomic support (Section 3.3.2), the TF family of its proximal gene (Section 3.3.3), and positional overlap with stress-associated MQTLs (Section 3.3.4). We additionally included functional annotations of proximal genes from MaizeGDB^6^ and the TF families represented by significant FIMO (Find Individual Motif Occurrences) [88] motif hits (FDR-adjusted *q* ≤ 0.05), identified within both the full ±3 kb sequence window surrounding the locus and the corresponding model-emphasized high-attention regions.

Considering the union of loci ranked within the top 10% by PT and PT+DoRA, the following locus- and gene-sharing patterns were observed across stress conditions:

- Although no individual SNP was shared across heat, drought, and combined heat-drought conditions, distinct prioritized SNPs within the LD half-decay distance (±16 kb) mapped to 13 proximal genes shared across all three conditions: *Zm00001eb019890, Zm00001eb026810, Zm00001eb042130, Zm00001eb141550, Zm00001eb141560, Zm00001eb203860, Zm00001eb204540, Zm00001eb206500, Zm00001eb301330, Zm00001eb304880, Zm00001eb372940, Zm00001eb416900,* and *Zm00001eb430490*.
- 32 prioritized SNPs were shared between heat and drought conditions. Of these, 9 SNPs were proximal to genes with transcriptomic support under both stresses: S3_6857744 (*Zm00001eb121370*), S4_210402356 (*Zm00001eb200830*), S5_34311892 and S5_34311958 (*Zm00001eb222730*), S6_127693439 (*Zm00001eb280520*), S6_128561728 (*intergenic: Zm00001eb280620-Zm00001eb280640*) S7_22513504 (*Zm00001eb303780*), S10_85891397 (*Zm00001eb416980*), and S10_108106279 (*intergenic: Zm00001eb420480-Zm00001eb420500*).

Moreover, 109 heat-, 57 drought-, and 45 combined heat-drought loci were prioritized by both methods, representing *the intersection of the top 10% loci identified by PT and PT+DoRA*. Among the heat- and drought-associated intersections, 50 heat loci (6 intergenic) and 19 drought loci (2 intergenic), had proximal genes overlapping DEGs in at least one transcriptomic study (Section 3.3.3), *providing convergent support across prioritization methods and transcriptomic evidence and making them highest confidence candidates (Table S1). Among these 69 loci, chromosomes 1, 5, and 4 contained the largest numbers of loci (11, 11, and 10, respectively)*.

Notable examples included the heat-responsive **GRAS** TF locus S1_159344889 (*Zm00001eb029390*) and several drought-associated candidates: the **G2-like** TF locus S4_36423454 (*Zm00001eb172880*), S4_177075897 within MQTL4.4 [77] (Zm00001eb191640), S4_238261570 within MQTL4.2 (*Zm00001eb205110*), and the intergenic locus S5_49834895 within MQTL5.3 (*Zm00001eb225310*-*Zm00001eb225320*). Additional prioritized loci on other chromosomes were proximal to genes encoding TFs from families, including **OVATE** (S3_222804153; *Zm00001eb159380*), **NAC** (S9_152472387; Zm00001eb399930), **Homeobox** (S10_85891397; upstream of Zm00001eb416980), and **WRKY** (S10_127270966; 5′ UTR of Zm00001eb424710).

For combined heat-drought stress, functional support was more limited: of the 45 loci prioritized by both methods, only S4_239877985 (3′UTR, Zm00001eb205480) located within an MQTL (MQTL4.2). Six additional loci were proximal to TF-encoding genes: S3_160670857 (Zm00001eb142960) and S3_199932499 (Zm00001eb152320–Zm00001eb152350, intergenic), both TCP; S1_284021511 (Zm00001eb057560, MADS); S3_180347114 (Zm00001eb146650, AP2/ERF-ERF); S4_244193607 (Zm00001eb207010, SBP); and S6_125016087 (Zm00001eb279900, LIM).

## 4. Discussion

Our results show that stress-conditioned adaptation of AgroNT reshaped sequence representations according to environmental context and produced biologically informative signals for prioritizing putative stress-responsive genomic regions. *Stress-specific prompt tokens with PT served as the primary conditioning mechanism, while LoRA and DoRA further improved embedding separation when combined with PT.* Notably, supervised contrastive fine-tuning was applied to the final-layer stress-token embeddings, whereas downstream prioritization relied on the DNA-token representations. The resulting stress-induced changes in the DNA-token representations indicate that environmental conditioning propagated through the transformer layers and modulated the sequence representations through interactions with the learned stress context.

However, improved separation of environmental contexts in the sequence embedding space did not translate uniformly into stronger biological support across downstream analyses. Among the adaptation strategies, PT+DoRA achieved the highest silhouette score and showed modest improvements over PT in DEG overlap, transcriptional-response patterns, proximal-gene TF-family characterization, and heat-associated MQTL overlap, whereas PT showed stronger enrichment for drought-associated MQTL overlap. These results suggest that the strategies captured complementary sequence features and that embedding separation alone does not fully predict downstream biological relevance.

Sequence-level motif enrichment identified abiotic stress-related signatures under all strategies. PT+LoRA additionally recovered larger motif sets under drought and combined heat-drought stress; however, these differences should be interpreted cautiously because the comparisons were not GC-matched. Score ablation suggested that combining embedding shift with attention Gini helped mitigate some of the compositional biases observed with the individual score components. GC-matched attention-guided analysis further showed that model-emphasized regions were enriched for known plant stress-regulatory motifs. Nevertheless, transformer attention should be interpreted as a model-derived prioritization signal rather than as direct evidence of TF binding.

Moreover, the prioritized loci overlapped only partially across the stress conditions, suggesting that combined-stress prioritization identified a largely distinct set of candidate loci rather than recapitulating heat-or drought-specific signals. Nonetheless, several combined stress loci overlapped heat-and drought-associated MQTLs, indicating regional convergence across stress conditions and suggesting that different variants located within broader stress-associated genomic intervals may be prioritized under different environment contexts. However, combined-stress analysis was constrained by the smaller number of year-location combinations available (Table 1), which reduced GWAS power for identifying initial candidate loci, and by the absence of directly comparable transcriptomic datasets for combined heat-drought stress.

Additionally, functional analysis of the loci prioritized by both PT and PT+DoRA (Supplementary File 1) revealed distinct biological processes under each condition, reflecting stress-specific adaptive mechanisms.

- Under heat stress, prioritized loci were proximal to genes encoding GRAS (S1_159344889), NAM (S9_152472387), and WRKY (S10_127270966) TFs, highlighting their potential roles in regulating heat-responsive pathways [89–91]. Several additional genes were associated with carbohydrate and energy metabolism, including a triose-phosphate transporter (S8_180898896), glycosyl hydrolase (S10_143606406), NAD-dependent glycerol-3-phosphate dehydrogenase (S7_22148622), thioredoxin (S4_7453738), a component associated with the pentose phosphate pathway (S3_156069459), and phosphoenolpyruvate synthase (S8_6796272). Transport-related genes, including K⁺ transporters (S2_9454374, S2_9456962), were also identified, suggesting potential roles for ion homeostasis and membrane transport in maintaining cellular function and physiological stability under elevated temperatures [92]. Collectively, these candidates suggest that heat-stress adaptation involves coordinated transcriptional regulation together with adjustments in carbon metabolism, energy balance, cellular redox homeostasis, and ion transport [93].
- Under drought stress, the prioritized loci were proximal to genes encoding MYB TFs (S4_36423454, S8_162896919, S10_139832700), which regulate diverse drought-responsive processes [94]. Genes involved in jasmonate signaling (12-oxo-phytodienoic acid reductase, S3_71244297), and flavonoid metabolism (dihydroflavonol-4-reductase, S5_49834895), were also prioritized, suggesting potential roles in stress signaling and antioxidant protection during drought.
- Under combined heat-drought stress, the prioritized loci were predominantly proximal to genes associated with lipid metabolism, including diacylglycerol acyltransferase (S3_68394121), diacylglycerol kinase (S10_19686195), and enoyl-CoA hydratase/isomerase (S10_54491391), suggesting potential roles in lipid remodeling, membrane stability, lipid signaling, and energy metabolism [95,96]. The prioritization of glutelin-2 (S7_125438681) and an ovule-associated protein (S10_33293936) may further indicate alterations in storage, resource allocation, and reproductive processes under combined stress.

Several high-confidence loci, prioritized by both PT and PT+DoRA with supporting transcriptomic evidence (Table S1, Section 3.3.6), were proximal to genes belonging to TF families with established roles in abiotic stress responses. **GRAS** TFs contribute to heat and drought resilience through gibberellin signaling, root development, and maintenance of cellular homeostasis [97,98], supporting the potential relevance of GRAS-associated locus identified here. **G2-like (GOLDEN2-LIKE; GLK)** factors regulate chloroplast development, photosynthetic capacity, and carbon assimilation during environmental stress [99]. Functional studies further support their role in stress tolerance: **SlGLK16** acts as a positive regulator of drought tolerance in tomato [100], while **TaG2-like** genes in wheat exhibit tissue-specific expression and responsiveness to abiotic stresses, including cold, ABA, salinity, and osmotic stress, suggesting conserved roles in stress adaptation [101]. **NAC** factors are central regulators of drought tolerance through ABA signaling, reactive oxygen species (ROS) scavenging, osmotic adjustment, and stress-induced senescence [102,103]. For example, **SNAC1** regulates the downstream phosphatase gene **OsPP18**, which contributes to drought and oxidative stress tolerance in rice by maintaining ROS homeostasis through an ABA-independent pathway [104]. Likewise, overexpression of **HvSNAC1** significantly improves drought tolerance in barley [105], highlighting the conserved role of NAC TFs in cereal stress adaptation. **WRKY** factors function as major regulatory hubs integrating ABA signaling, antioxidant defense, and transcriptional responses to drought and heat stress [106]. In maize, **ZmWRKY40** and **ZmWRKY106** positively regulate abiotic stress tolerance through activation of stress-responsive genes [91,107], while **GmWRKY27** enhances drought and salt tolerance in soybean by interacting with other TFs to regulate downstream stress-response pathways[108], suggesting that WRKY-mediated regulatory networks are broadly conserved across plant species. Although **OVATE family proteins (OFPs)** are less directly characterized in abiotic stress responses, their roles in brassinosteroid- and gibberellin-dependent regulation of plant architecture and cell elongation suggest a possible contribution through growth and morphological plasticity under adverse conditions [109]. **HD-ZIP factors** likewise integrate developmental and environmental signaling and have been implicated in ABA-mediated responses, water-deficit tolerance, and drought-responsive gene regulation [85,110].

Further, we analyzed the functional involvement genes proximal to loci prioritized by PT under drought stress, as this adaptation strategy showed the strongest enrichment for overlap with drought-associated MQTLs (Section 3.3.4). These genes were predominantly associated with primary and secondary metabolism, particularly pathways involved in carbohydrate metabolism, including glycosyl hydrolases (S2_237427328 and S1_33114289) and transferases (S3_3915406 and S9_3574078), which are important for carbohydrate remodeling and osmotic adjustment [111]. Additional candidate genes were linked to energy metabolism and the biosynthesis of secondary metabolites, including vitamin B12-related metabolism (S9_12406501 and S9_12406441), terpene biosynthesis (S9_13092307), and flavonoid biosynthesis through a 2OG-Fe(II) oxygenase superfamily gene (S5_3107066). These metabolic pathways suggest potential roles in energy redistribution, osmoprotection, antioxidant defense, and the accumulation of protective metabolites that may contribute to drought tolerance [112].

## 5. Conclusion

The proposed environment-aware DNA-LLM framework introduces an explicit mechanism for incorporating environmental stress context into a plant GFM through stress-specific prompt tokens and parameter-efficient adaptation. Unlike pre-trained GFMs that generate a single representation for a given genomic region, the framework produces stress-conditioned representations of the same DNA sequence under heat, drought, combined heat-drought, and control conditions. Building on these representations, we developed a prioritization strategy that integrates stress-induced representation shifts relative to the control condition with stress-token attention concentration to rank putative stress-responsive genomic regions.

Applied to maize grain yield, the learned stress-conditioned representations generalized across phenotypes without further updating the model parameters, as demonstrated by UMAP and silhouette-score analysis, indicating that the learned environmental context was not restricted to a single phenotype. The framework captured sequence-contextual signals complementary to conventional GWAS-based statistical association. The prioritized regions were further supported by overlap with stress-responsive DEGs and stress-associated MQTLs, regulatory motif analyses, and functional interpretation of proximal genes. Collectively, the prioritized loci and proximal genes represent biologically plausible candidates for plant stress adaptation, encompassing transcriptional regulation, stress signaling, redox and energy metabolism, and lipid remodeling. Their stress-specific patterns suggest that plants may employ distinct but interconnected molecular mechanisms to cope with heat, drought, and their combined effects.

Overall, these findings establish a methodological framework for integrating environmental context directly into GFMs and demonstrate its potential to connect sequence-level representation learning with genotype-by-environment analysis. More broadly, the proposed approach provides a foundation for leveraging AI in environment-aware genomic prioritization and could be extended to other crops, traits, and environmental conditions for crop improvement.

Several limitations should also be considered. The framework prioritizes GWAS-derived candidate regions but does not establish causal variants, regulatory elements, or target genes. LD-based gene assignment and MQTL overlap provide positional support rather than causal confirmation, and motif enrichment does not establish stress-specific regulatory activity *in vivo*. Sequence modeling was also restricted to ±3 kb around each locus by AgroNT’s input-length constraint, which is smaller than the LD half-decay distance in this panel (∼16 kb). Therefore, the model captures local sequence context but not the full genomic interval considered in LD-based gene assignment.

Future studies integrating chromatin-accessibility profiles, eQTLs, allele-specific expression, and experimental perturbation will be important for strengthening causal interpretation. Broader LD-based genomic context could also be explored through hierarchical encoding of multiple AgroNT-compatible windows. Other transformer-based GFMs capable of modeling longer genomic contexts, such as the Arabidopsis-trained GENA-LM (36 kb) [47], could further improve stress-conditioned genomic representations and locus prioritization while enabling assessment of cross-species transferability, although scaling to longer contexts requires efficiency mechanisms such as sparse attention. Non-transformer architectures (e.g., Evo 2 [52]) support substantially longer genomic contexts, extending to approximately 1 Mb. However, they do not provide transformer-style attention patterns; therefore, adapting the proposed framework would require alternative measures of local sequence importance for incorporation into the priority score.

## Acknowledgements

We thank Eleanor Carr for providing the code used for the GWAS analysis and Dr. Rajneesh Singhal for helpful discussions on biological concepts relevant to this study.

## Footnotes

1 https://www.genomes2fields.org/home/

2 https://github.com/iPRoBe-lab/Environment_Aware_DNA_LLM.git

3 https://maizemine.rnet.missouri.edu/maizemine/

4 https://grassius.org/species/Maize

5 https://maizegdb.org

6 https://download.maizegdb.org/GeneFunction_and_Expression/ProtNLM/Uniprot_ProtNLM_maize_v5.txt

