## Supplementary Materials for "Environment-Aware DNA Language Model for Stress-Responsive Genomic Prioritization in Maize"

Supplementary Figures

**Fig. S2. Manhattan plots of GWAS results for grain yield under different stress conditions.** The Bonferroni threshold was defined as 0.05 divided by the number of tested markers, and the FDR threshold was estimated using the Benjamini–Hochberg procedure. A permissive threshold of p ≤ 0.05 was used to retain a broader set of stress-associated candidate loci for sequence-based modeling.

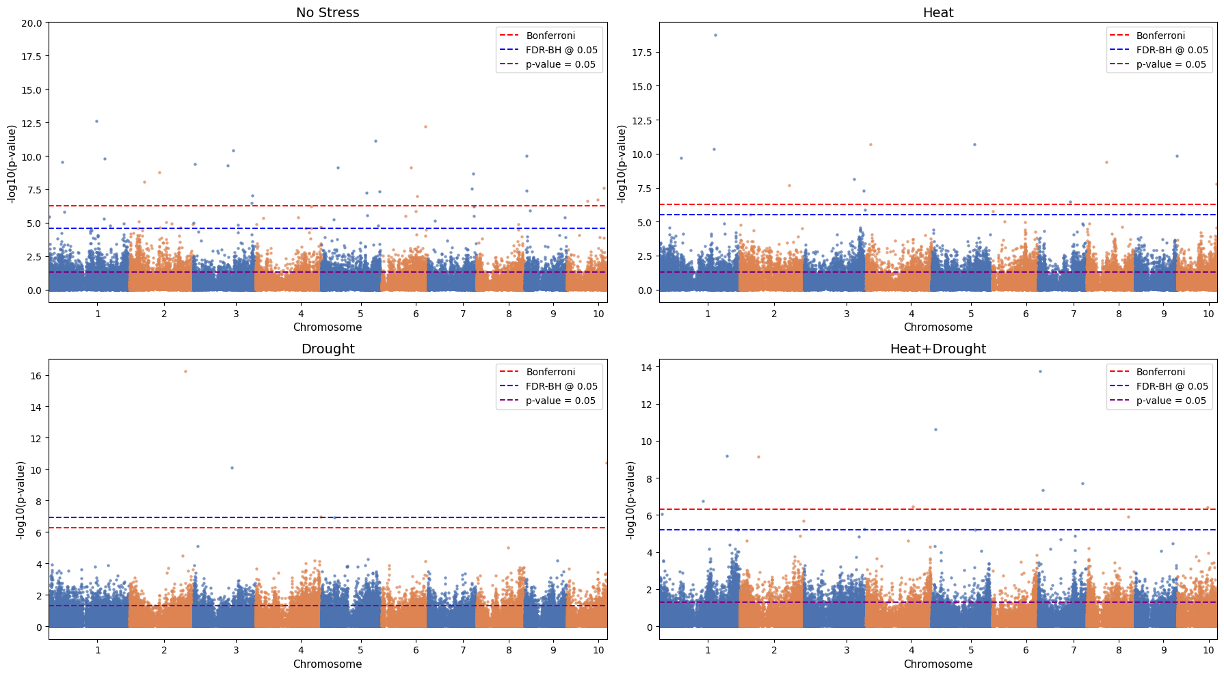

**Fig. S1.** LD decay across the G2F unpruned maize hybrid panel (437,214 SNPs). Mean pairwise squared genotype correlation (r²) between SNPs, as a function of physical distance, with the fitted Hill–Weir decay curve (red). The estimated background LD plateau was r² ≈ 0.16 (grey dotted line), and the half-decay distance of 15.7 kb (purple dashed line); ±32 kb (twice the half-decay distance) is shown as a conservative bound (orange dashed line).

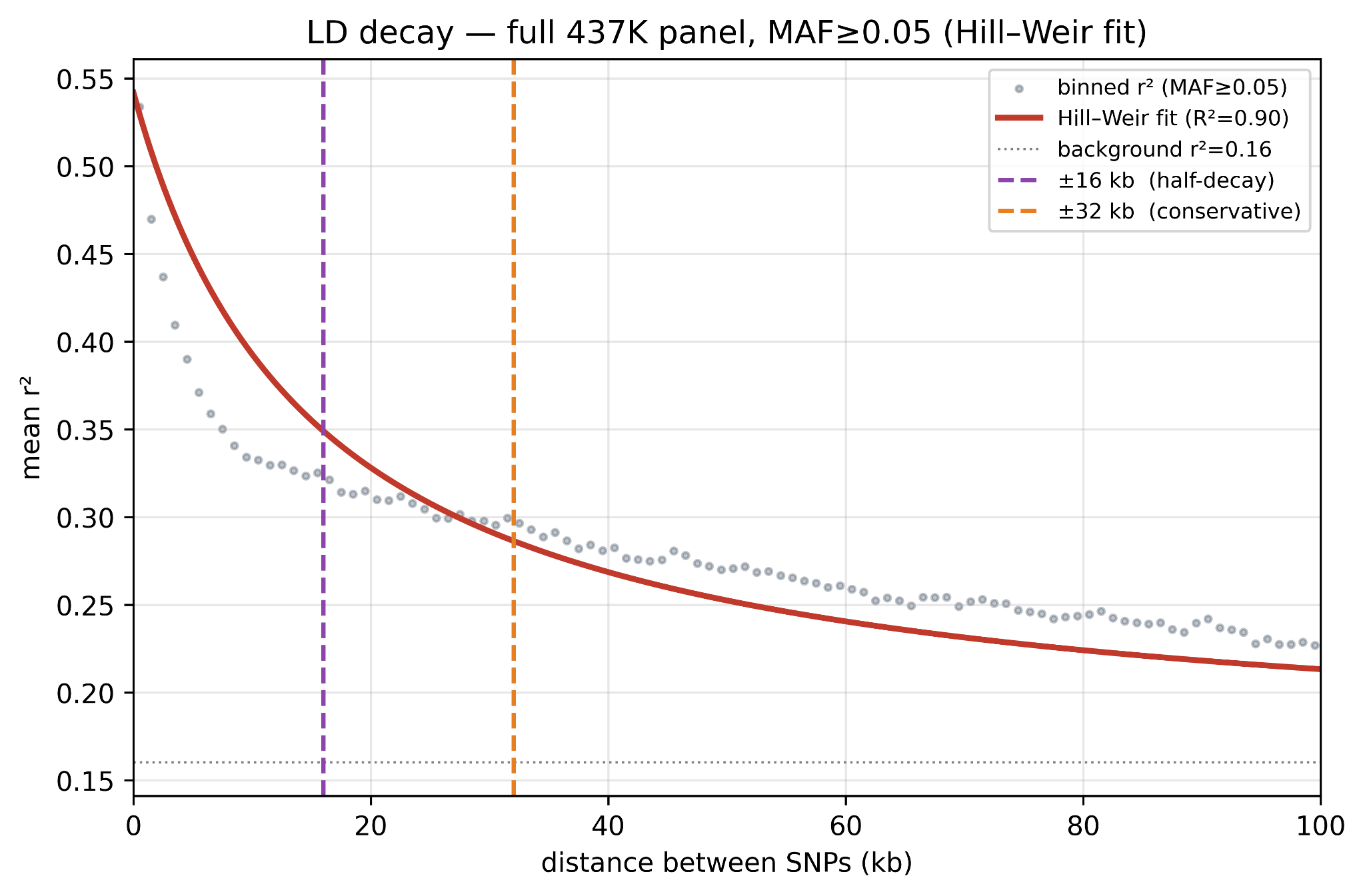

**Fig. S3. Cross-phenotype generalization of stress-conditioned sequence embeddings.** UMAP of flowering trait-associated genomic sequences generated by the stress-conditioned AgroNT model after prompt tuning (PT) on yield-associated sequences, without further fine-tuning. Input sequences correspond to genomic regions spanning ±3 kb around GWAS-identified loci (p ≤ 0.05).

(a) Anthesis

(b) Silking

(c) ASI

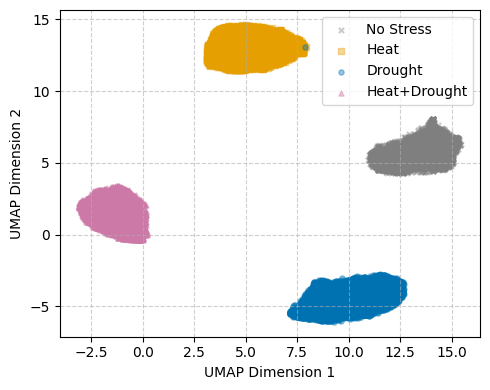

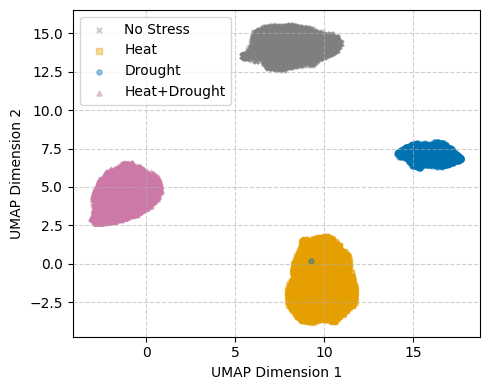

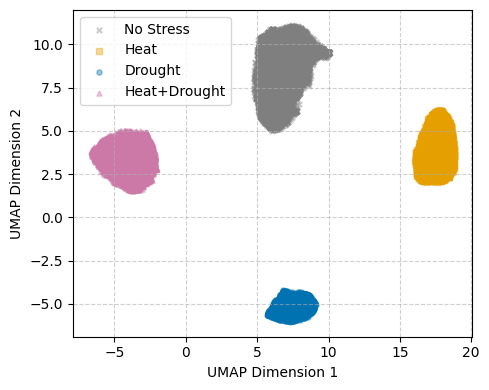

**Fig. S4. In silico contextual perturbation by counterfactual prompting.** UMAP of sequence embeddings for yield-associated loci (p ≤ 0.05) under the original control condition, the original stress-conditioned input, and the counterfactual input generated by replacing the stress prompt with <NO_STRESS> while keeping the nucleotide sequence fixed. Counterfactual embeddings shifted toward the control cluster, indicating that the prompt context contributes substantially to stress-specific embedding structure.

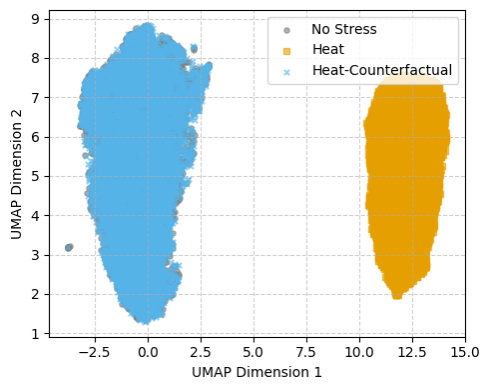

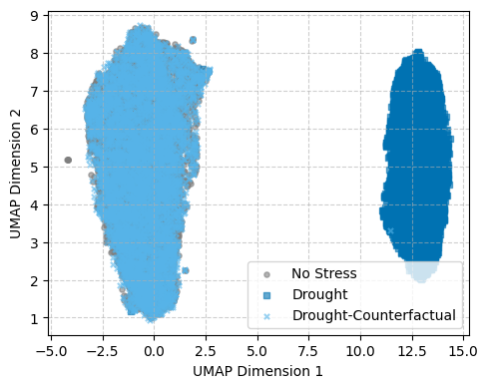

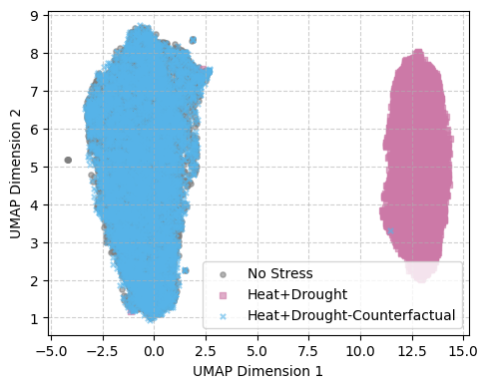

(a) Heat

(b) Drought

(c) Heat+Drought

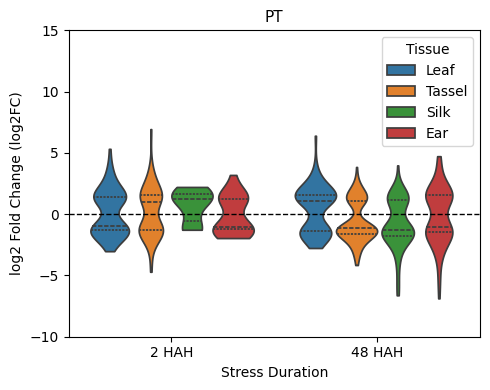

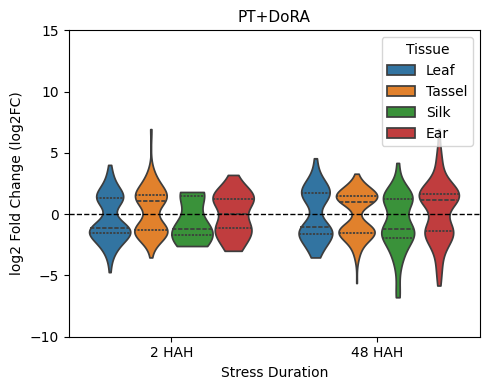

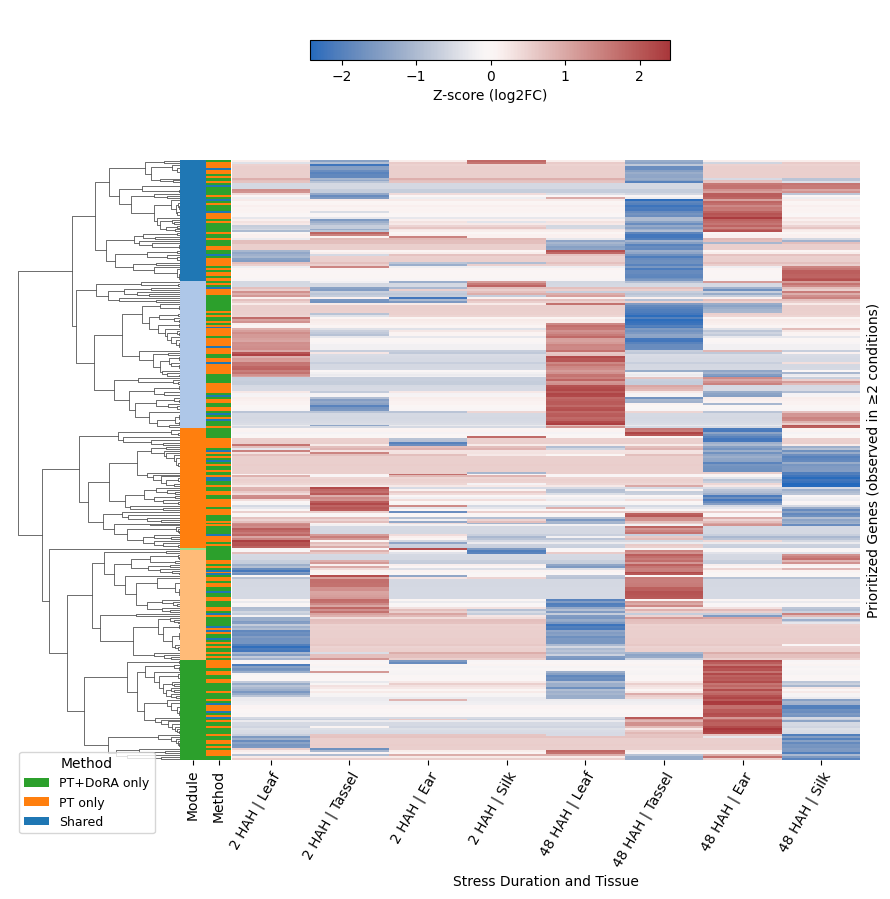

**Fig. S5. Expression profiles and gene-level clustering of heat-prioritized proximal genes overlapping with DEGs reported by He et al. (2019).** (a–b) Violin plots summarizing the distribution of log₂ fold-change (log₂FC) values across tissues and stress durations. (c) Heatmap of row-wise z-score-normalized log₂FC values for genes observed in at least two tissue–duration conditions. Genes were hierarchically clustered using correlation distance and average linkage, with modules defined at a distance threshold of 0.8. Row annotations indicate module membership and prioritization category (PT only, PT+DoRA only, or shared).

Supplementary Tables

**Table S1.** Prioritized loci identified by both PT and PT+DoRA whose proximal genes overlapped with DEGs reported in at least one of the transcriptomic studies under heat stress [66–71]

| **Sl. No.** | **SNP** | **Stress Factor** | **Annotation** | **Gene ID** | **TF Family** | **MQTL ID [77]** | **High-Attention Motif Family** | |
| --- | --- | --- | --- | --- | --- | --- | --- | --- |
|  |  |  |  |  |  |  | **PT** | **PT+DoRA** |
| 1 | S1_60310410 | Heat | Downstream | Zm00001eb016940 |  |  | DOF | DOF; ERF/DREB |
| 2 | S1_62475593 | Heat | Upstream | Zm00001eb017450 |  |  | BBR/BPC; DOF | DOF |
| 3 | S1_98821878 | Heat | Intergenic | Zm00001eb024230-Zm00001eb024240 |  |  | BBR/BPC; C2H2; DOF | C2H2; DOF; ERF/DREB |
| 4 | S1_159344889 | Heat | 5'-UTR | Zm00001eb029390 | GRAS |  | DOF; PLINC | DOF; ERF/DREB; LBD |
| 5 | S1_183679792 | Drought | Exon | Zm00001eb032790 |  |  | DOF; ERF/DREB | BBR/BPC; DOF; ERF/DREB; LBD |
| 6 | S1_194954652 | Heat | Exon | Zm00001eb035700 |  |  | BBR/BPC; DOF; ERF/DREB | BBR/BPC; DOF; ERF/DREB; LBD |
| 7 | S1_200450010 | Heat | Exon | Zm00001eb037120 |  |  | DOF | DOF; ERF/DREB |
| 8 | S1_229360995 | Heat | Intron | Zm00001eb043650 |  |  | BBR/BPC; C2H2; DOF | BBR/BPC; C2H2; DOF; ERF/DREB; LBD |
| 9 | S1_277193426 | Drought | Exon | Zm00001eb055750 |  |  | CPP; DOF; ERF/DREB | CPP; ERF/DREB |
| 10 | S1_292009202 | Heat | Exon | Zm00001eb059900 |  |  | DOF | DOF; ERF/DREB |
| 11 | S1_307035004 | Heat | Exon | Zm00001eb065160 |  |  | BBR/BPC | ERF/DREB |
| 12 | S2_10434135 | Heat | Upstream | Zm00001eb070760 |  |  | BBR/BPC; DOF; ERF/DREB; LBD | BBR/BPC; C2H2; DOF; ERF/DREB; LBD |
| 13 | S2_168132731 | Drought | Upstream | Zm00001eb095660 |  |  | BBR/BPC; DOF; ERF/DREB; LBD | ERF/DREB; LBD |
| 14 | S2_74560607 | Heat | Upstream | Zm00001eb085610 |  |  | BBR/BPC; C2H2; DOF; MIKC | BBR/BPC; C2H2; DOF; ERF/DREB; LBD; MIKC |
| 15 | S2_103377926 | Heat | Intergenic | Zm00001eb087830-Zm00001eb087840 |  |  | BBR/BPC; CPP; DOF | BBR/BPC; C2H2; CPP; DOF; ERF/DREB; LBD |
| 16 | S3_130587307 | Drought | 3'-UTR | Zm00001eb137020 |  |  | BBR/BPC; DOF | BBR/BPC; DOF; ERF/DREB; LBD |
| 17 | S3_145521256 | Heat | Exon | Zm00001eb139660 |  |  | BBR/BPC; C2H2; DOF; ERF/DREB; LBD | DOF; ERF/DREB; LBD |
| 18 | S3_156069459 | Heat | Exon | Zm00001eb141930 |  |  | DOF | DOF; ERF/DREB |
| 19 | S3_173101064 | Heat | 3'-UTR | Zm00001eb144910 |  |  | DOF | DOF; ERF/DREB |
| 20 | S3_194887297 | Heat | Upstream | Zm00001eb151170 |  |  | BBR/BPC; DOF | BBR/BPC; C2H2; DOF; ERF/DREB; LBD |
| 21 | S3_217000251 | Heat | Intergenic | Zm00001eb157600-Zm00001eb157610 |  |  | BBR/BPC; DOF | BBR/BPC; DOF; ERF/DREB |
| 22 | S3_222804153 | Heat | Downstream | Zm00001eb159380 | OVATE |  | BBR/BPC; DOF | BBR/BPC; DOF; ERF/DREB; LBD |
| 23 | S3_230262078 | Heat | 5'-UTR | Zm00001eb161860 |  |  | DOF | DOF; ERF/DREB |
| 24 | S4_14489145 | Drought | Exon | Zm00001eb168550 |  |  | DOF; ERF/DREB; LBD | DOF; ERF/DREB; LBD |
| 25 | S4_36423454 | Drought | 5'-UTR | Zm00001eb172880 | G2-like |  | BBR/BPC; DOF; ERF/DREB | AP2; ERF/DREB |
| 26 | S4_6690441 | Heat | Exon | Zm00001eb166980 |  |  | DOF | BBR/BPC; DOF; ERF/DREB |
| 27 | S4_163538058 | Drought | Exon | Zm00001eb187980 |  |  | DOF; ERF/DREB; LBD | DOF; ERF/DREB; LBD |
| 28 | S4_177075897 | Drought | Upstream | Zm00001eb191640 |  | MQTL4.4 | BBR/BPC; DOF; ERF/DREB; LBD | ERF/DREB; LBD |
| 29 | S4_207676296 | Heat | Intron | Zm00001eb200400 |  |  | DOF | BBR/BPC; DOF; ERF/DREB; LBD |
| 30 | S4_228694080 | Heat | Intergenic | Zm00001eb203570-Zm00001eb203600 |  |  | BBR/BPC; C2H2; DOF | BBR/BPC; C2H2; DOF; ERF/DREB; LBD |
| 31 | S4_238261570 | Drought | Upstream | Zm00001eb205110 |  | MQTL4.2 | BBR/BPC; C2H2; CPP; DOF; ERF/DREB; LBD | BBR/BPC; C2H2; DOF; ERF/DREB; LBD |
| 32 | S4_243087461 | Heat | Upstream | Zm00001eb206400 |  |  | BBR/BPC; CPP; DOF; ERF/DREB; LBD | BBR/BPC; CPP; DOF; ERF/DREB; LBD |
| 33 | S4_248492375 | Heat | Upstream | Zm00001eb209360 |  |  | DOF; MIKC | DOF; ERF/DREB |
| 37 | S5_4270288 | Heat | Upstream | Zm00001eb212800 |  |  | BBR/BPC; CPP; DOF | BBR/BPC; C2H2; DOF; ERF/DREB |
| 38 | S5_4270349 | Heat | Upstream | Zm00001eb212800 |  |  | BBR/BPC; DOF; ERF/DREB | C2H2; ERF/DREB |
| 39 | S5_4756985 | Drought | Exon | Zm00001eb213120 |  |  | BBR/BPC; C2H2; DOF; ERF/DREB; LBD | BBR/BPC; C2H2; ERF/DREB |
| 34 | S5_34311892 | Heat | Exon | Zm00001eb222730 |  |  | BBR/BPC; DOF; ERF/DREB | BBR/BPC; DOF; ERF/DREB; LBD |
| 40 | S5_34311958 | Heat | Exon | Zm00001eb222730 |  |  | BBR/BPC; DOF; ERF/DREB | BBR/BPC; DOF; ERF/DREB; LBD |
| 35 | S5_77148826 | Heat | Intron | Zm00001eb231370 |  |  | DOF | ERF/DREB |
| 36 | S5_85227512 | Drought | Upstream | Zm00001eb232850 |  |  | BBR/BPC; C2H2; DOF; ERF/DREB; LBD | BBR/BPC; C2H2; ERF/DREB; LBD |
| 41 | S5_49834895 | Drought | Intergenic | Zm00001eb225310-Zm00001eb225320 |  | MQTL5.3 | BBR/BPC; C2H2; DOF; ERF/DREB | BBR/BPC; C2H2; ERF/DREB |
| 42 | S5_163146528 | Drought | Downstream | Zm00001eb240930 |  |  | BBR/BPC; C2H2; DOF; ERF/DREB; LBD | BBR/BPC; DOF; ERF/DREB; LBD |
| 43 | S5_171984360 | Drought | 5'-UTR | Zm00001eb242690 |  |  | BBR/BPC; C2H2; DOF; ERF/DREB; LBD | BBR/BPC; C2H2; DOF; ERF/DREB |
| 44 | S5_194815084 | Heat | Exon | Zm00001eb249020 |  |  | DOF | BBR/BPC; ERF/DREB; LBD |
| 45 | S6_29498976 | Heat | Upstream | Zm00001eb264590 |  |  | BBR/BPC; C2H2; DOF | BBR/BPC; DOF; ERF/DREB |
| 46 | S6_47930491 | Heat | Downstream | Zm00001eb267110 |  |  | BBR/BPC; DOF | BBR/BPC; C2H2; DOF; ERF/DREB |
| 47 | S6_128561728 | Heat | Intergenic | Zm00001eb280620-Zm00001eb280640 |  |  | DOF | ERF/DREB |
| 48 | S6_129756497 | Heat | Upstream | Zm00001eb280810 |  |  | DOF | DOF; ERF/DREB; LBD |
| 49 | S6_147314912 | Heat | Exon | Zm00001eb285170 |  |  | DOF | BBR/BPC; C2H2; DOF; ERF/DREB |
| 50 | S6_166701426 | Heat | Upstream | Zm00001eb291190 |  |  | DOF | DOF; ERF/DREB |
| 51 | S6_166701854 | Heat | Upstream | Zm00001eb291190 |  |  | CPP; DOF | DOF; ERF/DREB |
| 52 | S6_172683650 | Heat | Exon | Zm00001eb294020 |  |  | BBR/BPC; DOF | BBR/BPC; DOF; ERF/DREB |
| 53 | S6_174244533 | Heat | Upstream | Zm00001eb294960 |  |  | CPP; DOF | DOF; ERF/DREB; LBD |
| 54 | S7_7730479 | Drought | Exon | Zm00001eb300730 |  |  | BBR/BPC; C2H2; DOF; ERF/DREB; LBD | BBR/BPC; C2H2; DOF; ERF/DREB |
| 55 | S7_10078891 | Drought | Exon | Zm00001eb301330 |  |  | BBR/BPC; DOF; ERF/DREB | BBR/BPC; DOF; ERF/DREB |
| 56 | S7_22148622 | Heat | Upstream | Zm00001eb303710 |  |  | BBR/BPC; DOF | BBR/BPC; DOF; ERF/DREB; LBD |
| 57 | S8_61171618 | Drought | Intergenic | Zm00001eb342080-Zm00001eb342090 |  |  | BBR/BPC; C2H2; DOF; ERF/DREB | BBR/BPC; DOF; ERF/DREB |
| 58 | S8_179592333 | Drought | Exon | Zm00001eb370260 |  |  | BBR/BPC; C2H2; DOF; ERF/DREB; LBD | BBR/BPC; DOF; ERF/DREB; LBD |
| 59 | S8_180898896 | Heat | Exon | Zm00001eb370980 |  |  | DOF | BBR/BPC; DOF; ERF/DREB; LBD |
| 60 | S9_152472387 | Heat | Downstream | Zm00001eb399930 | NAC |  | DOF; ERF/DREB | BBR/BPC; DOF; ERF/DREB |
| 61 | S10_28850137 | Heat | Intron | Zm00001eb410930 |  |  | BBR/BPC; DOF | BBR/BPC; DOF; ERF/DREB; LBD |
| 62 | S10_85891397 | Drought | Upstream | Zm00001eb416980 | Homeobox |  | BBR/BPC; C2H2; DOF; ERF/DREB; LBD | BBR/BPC; C2H2; DOF; ERF/DREB; LBD |
| 63 | S10_99565835 | Heat | Intergenic | Zm00001eb419330-Zm00001eb419350 |  |  | BBR/BPC; CPP; DOF | DOF |
| 64 | S10_127270966 | Heat | 5'-UTR | Zm00001eb424710 | WRKY |  | CPP; DOF | DOF; ERF/DREB |
| 65 | S10_142157971 | Heat | Exon | Zm00001eb429930 |  |  | DOF | DOF; ERF/DREB |
| 66 | S10_143614697 | Heat | Splice_site_region | Zm00001eb430480 |  |  | CPP; DOF | DOF; ERF/DREB |
| 67 | S10_144729792 | Heat | Exon | Zm00001eb430990 |  |  | DOF | BBR/BPC; DOF; ERF/DREB |
| 68 | S10_149269892 | Heat | Exon | Zm00001eb432990 | Orphans |  | DOF | DOF; ERF/DREB |
| 69 | S10_149717459 | Heat | Exon | Zm00001eb433330 |  |  | DOF; ERF/DREB | DOF; ERF/DREB; LBD |
